# Virus-Dependent Immune Conditioning of Tissue Microenvironments

**DOI:** 10.1101/2021.05.21.444548

**Authors:** Sizun Jiang, Chi Ngai Chan, Xavier Rovira-Clave, Han Chen, Yunhao Bai, Bokai Zhu, Erin McCaffrey, Noah F. Greenwald, Candace Liu, Graham L Barlow, Jason L. Weirather, John Paul Oliveria, Darci Philips, Nilanjan Mukherjee, Kathleen Busman-Sahay, Michael Nekorchuk, Margaret Terry, Skyler Younger, Marc Bosse, Janos Demeter, Yury Golstev, David Robert McIlwain, Michael Angelo, Jacob D. Estes, Garry P. Nolan

**Author notes:** Equal Contributions (Sizun Jiang, Chi Ngai Chan, Xavier Rovira-Clave). Senior Authors (Michael Angelo, Jacob D. Estes, Garry P. Nolan). Correspondence &.

## Abstract

A thorough understanding of complex spatial host-disease interactions *in situ* is necessary in order to develop effective preventative measures and therapeutic strategies. Here, we developed <u>P</u>rotein <u>A</u>nd <u>N</u>ucleic acid <u>IN</u> situ <u>I</u>maging (PANINI) and coupled it with Multiplexed Ion Beam Imaging (MIBI) to sensitively and simultaneously quantify DNA, RNA, and protein levels within the microenvironments of tissue compartments. The PANINI-MIBI approach was used to measure over 30 parameters simultaneously across large sections of archival lymphoid tissues from non-human primates that were healthy or infected with simian immunodeficiency virus (SIV), a model that accurately recapitulates human immunodeficiency virus infection (HIV). This enabled multiplexed dissection of cellular phenotypes, functional markers, viral DNA integration events, and viral RNA transcripts as resulting from viral infection. The results demonstrated immune coordination from an unexpected upregulation of IL10 in B cells in response to SIV infection that correlated with macrophage M2 polarization, thus conditioning a potential immunosuppressive environment that allows for viral production. This multiplexed imaging strategy also allowed characterization of the coordinated microenvironment around latently or actively infected cells to provide mechanistic insights into the process of viral latency. The spatial multi-modal framework presented here is applicable to deciphering tissue responses in other infectious diseases and tumor biology.

## Introduction

Since the beginning of the human immunodeficiency virus infection and acquired immunodeficiency syndrome (HIV/AIDS) pandemic in the 1980s, the majority of our knowledge of the biology and persistence of HIV-1 in humans and of its closely related cousin in non-human primates, the simian immunodeficiency virus (SIV), has come from studies of the peripheral blood compartment and the molecular biology of the virus within host cells *in vitro*. These experiments led to the development of the modern antiretroviral therapy (ART), which prevents the fatal progression to immunodeficiency in most patients (Hartman and Buckheit, 2012). Unfortunately, ART is not a cure for HIV/AIDS as viral rebound occurs in all but the rarest cases upon ART withdrawal. Furthermore, much of the viral replication and persistence during ART occurs within the lymphoid tissues and gastrointestinal tract (Chun et al., 1997; Estes et al., 2017; Haase, 1999). To understand the mechanisms and pathology of HIV persistence it is necessary to visualize the tissue microenvironments where the virus resides; however, technological barriers have limited our ability to phenotypically characterize and quantify the cellular components of viral tissue reservoirs.

Current approaches to study of the microenvironments of viral reservoirs include single-cell RNA-seq-based (Kazer et al., 2020) and flow-based (Baxter et al., 2017) methods that require cells to be taken out of their native tissue context. Complementary methods such as immunohistochemistry and *in situ* hybridization (ISH) technologies (Deleage et al., 2016; Estes et al., 2017) enable retention of the information in 2D space but are constrained by the low number of concurrently detectable features. Multiplexing markers on a tissue section using immunofluorescence microscopy is possible and routine but is generally limited by factors such as the spectral overlap of fluorophores and incompatible host species for primary antibodies. Recent advances in multiplexed imaging modalities, such as Multiplexed Ion Beam Imaging (MIBI) (Angelo et al., 2014; Keren et al., 2018), CO-Detection by indEXing (CODEX) (Goltsev et al., 2018; Schürch et al., 2020), Imaging Mass Cytometry (IMC) (Giesen et al., 2014), signal amplification by exchange reaction (SABER) (Kishi et al., 2019; Saka et al., 2019), and cyclic Immunofluorescence (cycIF) (Lin et al., 2015, 2018), which utilize iterative methods (CODEX, SABER and cycIF) or mass spectrometry (MIBI and IMC) to overcome these limitations. Highly multiplexed *in situ* detection of mRNA and protein epitopes has also been achieved with a branched-chain amplification method coupled with antibody-based detection (Schulz et al., 2018; Wang et al., 2012), but this procedure has been validated only for highly abundant RNA transcripts and requires a protease treatment step to increase RNA accessibility for detection sensitivity that interferes with robust protein epitope detection via antibodies (Schulz et al., 2018).

The ability to simultaneously detect nucleic acids present at low abundance, such as a single copy of the DNA resulting from a viral integration event, and protein molecules *in situ*, is paramount for enabling studies of viral infection. The low frequency of certain replication intermediates and need for high stringency hybridization requires a level of sensitivity that is not compatible with currently employed multiplexed imaging platforms. For example, the stripping buffers used in CODEX and the ethanol H_2_O desalting steps used in MIBI will disrupt most nucleic acid hybridizations. We reasoned that combining a customized branched-chain amplification method capable of DNA and RNA single-event detection with the covalent deposition of haptens would enable multiplexed imaging on various antibody-based platforms, including MIBI. By turning nucleic acid detection into an antibody “problem”, we could potentially overcome the limited sensitivity of ISH in tissues. For example, on the IMC and MIBI platforms, each oligonucleotide-based probe can only carry a maximum of 20 metal ions (Frei et al., 2016). In comparison, each antibody has a theoretical capacity for approximately 100 metal ions (Bendall et al., 2011; Han et al., 2018).

Here, we present an approach that we call <u>P</u>rotein <u>A</u>nd <u>N</u>ucleic acid <u>IN</u> situ <u>I</u>maging (PANINI) which when coupled to MIBI (PANINI-MIBI) provides 1) a highly sensitive custom branched-chain amplification method for nucleic acid targets with tyramide-based amplification, 2) an optimized antigen retrieval protocol that bypasses a protease treatment step and yet allows nucleic acid detection down to a single genomic event, and 3) antibody-based detection with resolution of down to 260 nm and single antibody sensitivity via the MIBI platform. Using formalin-fixed paraffin-embedded (FFPE) cell pellets and lymphoid tissues from SIV-infected and uninfected rhesus macaques, a system that appears to accurately reflect HIV infection in humans, we demonstrate that PANINI-MIBI is capable of simultaneous detection of single-integration events of SIV DNA (vDNA), SIV RNA transcripts (vRNA), and protein epitopes robustly on the same tissue section.

We utilized PANINI-MIBI to characterize in unprecedented detail the viral reservoir and immune responses within SIV-infected and uninfected control lymphoid tissues. The tissue immune responses to lentiviral infection were heterogenous, and phenotypically similar cells from infected animals and uninfected controls exhibited significantly different functions. For instance, IL10 expression was increased in B cells upon infection, thus promoting a presumed polarization of macrophages to an immunosuppressive M2 phenotype, which was correlated to a known conducive environment for SIV transcriptional activation. Characterization of the higher order structure around infected cells revealed differences during viral latency and active infection, enabling us to establish a model for how chronic SIV infection dampens the immune response and elucidate the hallmarks of host features that coordinate viral latency in tissue reservoirs. This work provides a framework for future multi-modal studies of the principles of host-pathogen interactions *in situ* using inactivated archival tissue samples.

## Results

### Development of PANINI

We designed the workflow of PANINI staining of tissue sections for subsequent analysis on the MIBI platform to be analogous to routine *in situ* hybridization and immunohistochemistry methods (Figure 1A). We first tested this approach using both immunofluorescence (IF) and MIBI on 3D8 and CEM FFPE cell pellets (Figure 1B). The 3D8 cell line was derived from a clonally expanded SIVmac316-infected CEM cell, containing a single copy of integrated SIV vDNA per cell (Nishimura et al., 2009). A single vDNA-positive punctate is expected within each 3D8 cell, and there is a 21-29% probability of capturing a positive nuclear event within 4-6-μm sections (Deleage et al., 2016). Quantification of vDNA-positive puncta in 3D8 cells from our IF and MIBI data was highly reproducible, and results from the two techniques aligned with each other and with previous studies (Figure 1C) (Deleage et al., 2016). This high concordance is indicative of the applicability of PANINI for sensitive, targeted detection of nucleic acids, down to a single SIV integration event, in FFPE archival tissue samples.

**Figure 1:**
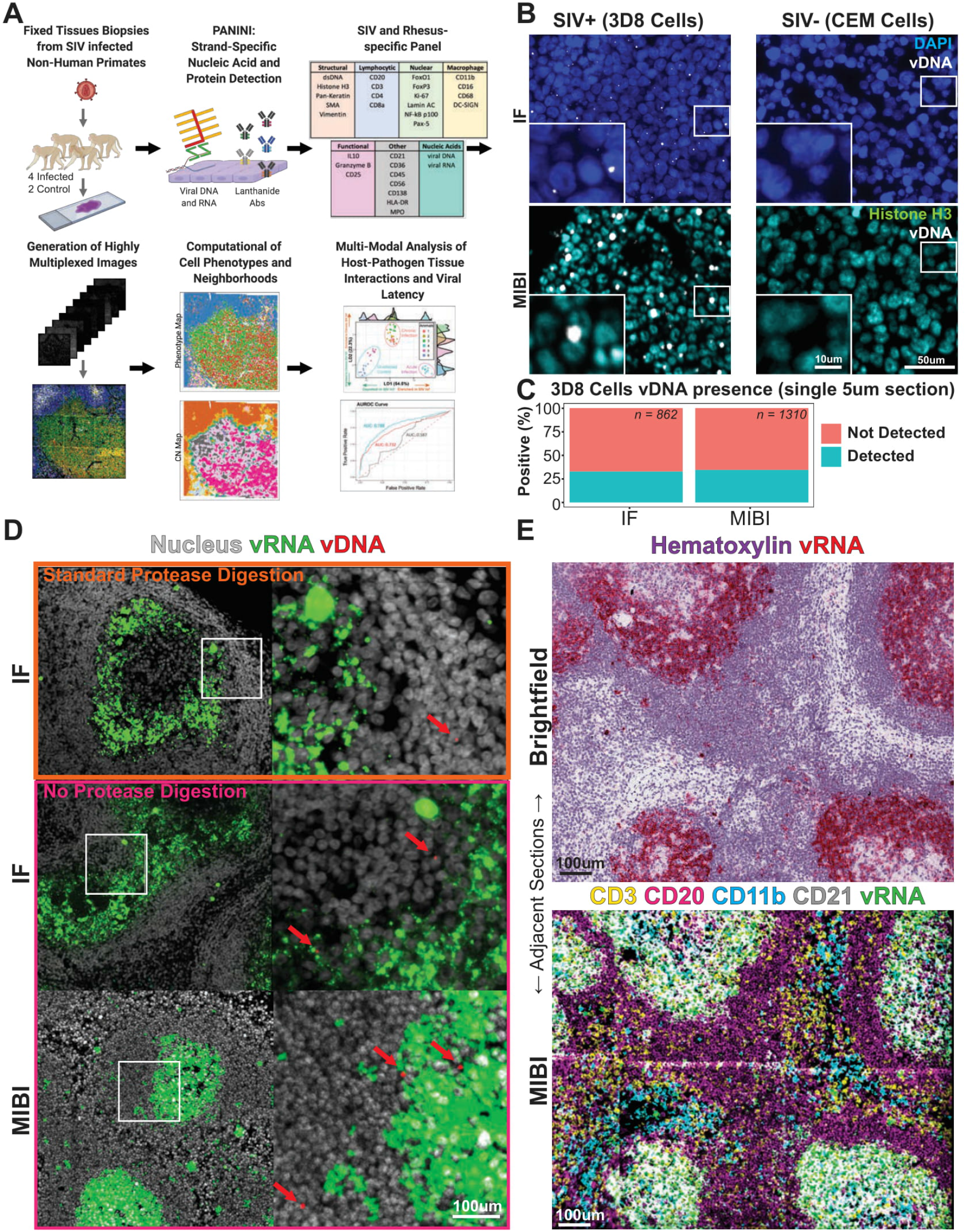
PANINI Enables Multiplexed, Strand-Specific Nucleic Acid and Protein Detection in Archival Tissues. **(A)** An overview of the experimental workflow and analytical framework for PANINI. In short, tissue autopsy sections from rhesus macaques were subject to the PANINI methodology, which couples nucleic acid amplification with antibody-based detection of both nucleic acid and protein targets. Multiplexed images were then acquired using MIBI and computationally analyzed for a high-resolution understanding of host-pathogen tissue interactions *in situ*. **(B)** Representative IF and MIBI images of positive control 3D8 cells and negative control CEMs. Nuclear stains, DAPI for IF and Histone H3 for MIBI are in blue and cyan respectively; vDNA is in white. **(C)** Quantification of 3D8 cells that are positive for vDNA signals in IF and MIBI images. **(D)** Representative images of SIV-positive lymph nodes subjected to standard protease digestion step (top) after epitope retrieval or no protease treatment (middle and bottom) after epitope retrieval. SIV vDNA (red) and vRNA (green) were detected using ISH followed by hapten deposition, and subsequently imaged using either IF or MIBI. Cells harboring integrated virus are indicated with red arrows. **(E)** Representative images of adjacent sections from an SIV-positive lymph node. Both slides underwent PANINI treatment. For the top section, a Fast Red chromogenic substrate was used for vRNA (red) and hematoxylin staining enabled brightfield visualization. The bottom section was stained with a MIBI-compatible protein cocktail. Markers shown here were selected to best delineate specific cell lineages: CD3 (T cells, yellow), CD20 (B cells, purple), CD11b (monocytes, blue), CD21 (B cells and FDCs, white), and SIV vRNA (green).

### Application of PANINI with MIBI

The protease digestion step used in various ISH assays, including the popular RNAscope and its related DNAscope (Deleage et al., 2016; Wang et al., 2012), is required to increase the accessibility of target nucleic acids through disruption of the packed architecture of tissue matrixes and degradation of nucleic acid-binding proteins (Yang et al., 1998). We found that a pH 9 antigen retrieval step enabled detection of vDNA and vRNA in FFPE lymph node sections from an SIV-infected rhesus macaque without the need for protease digestion (Figure 1D, top). These results are in line with previous observations that treatment of tissues with base can facilitate DNA recovery from FFPE tissue samples (Shi et al., 2004). Similar results were obtained by IF and MIBI without protease digestion (Figure 1, middle and bottom).

To better analyze the dynamic immune response to chronic SIV infection, we validated and applied a 33-marker panel that included probes to vDNA and vRNA (Figure S1A) across lymphoid tissues from four SIV-infected and two uninfected rhesus macaques, resulting in approximately 470,000 spatially resolved cells. The staining specificity was thoroughly assessed (Figure S1B). Analyses of adjacent sections of a lymph node from an SIV-infected rhesus macaque were subject to standard single-plex RNAscope ISH (Figure 1E, top) or PANINI-MIBI (Figure 1E, bottom, and Figure S1C), demonstrating that the latter captures viral events (SIV vRNA) and tissue morphology while significantly expanding upon the number of markers that can be simultaneously assessed. This is exemplified by CD3 for T cells, CD20 for B cells, CD11b for monocytes, and CD21 for B cells and follicular dendritic cells (FDCs) (Figure 1E, bottom). The ability to visualize multiple lineage-specific markers simultaneously enables both cross-validation and detailed phenotyping, such as for regulatory T cells (Tregs; Figure S1D), granzyme B^+^ CD8^+^ T cells (Figure S1E), B cells and FDCs (Figure S1F), and M1/M2 macrophages (Figure S1G).

### Scalable Automated Cell Segmentation and Annotation

Accurate cell segmentation methods are required to confidently extract single-cell feature information from multiplexed tissue images (Hollandi et al., 2020; Moen et al., 2019; Valen et al., 2016). We used the top-in-class Mesmer, a DeepCell-based segmentation method for feature extraction and FlowSOM for subsequent cell type identification utilizing self-organizing maps (Figure 2A) (Gassen et al., 2015; Greenwald et al., 2021). We identified 14 distinct immune and structural cell types, with the expected associated lineage-specific marker expression (Figure 2B). Visual inspection of the MIBI multiplexed images (Figure 2C) and their paired spatial phenotype maps confirmed accurate celltype annotation (Figure S2A). We orthogonally performed immunofluorescent immunohistochemistry on three consecutive sections juxtaposed to the original PANINI-MIBI analyzed section, which confirmed specificity and scalability of the unsupervised cell annotation methodology (Figure S2B). Prominent tissue features, such as B cell follicles, T cell zones, and the macrophage-rich medullary sinus, were visible on both the MIBI images (Figure 2C) and phenotype maps (Figure 2D and S3), further confirming the robustness of the cell segmentation and annotation methodology.

**Figure 2:**
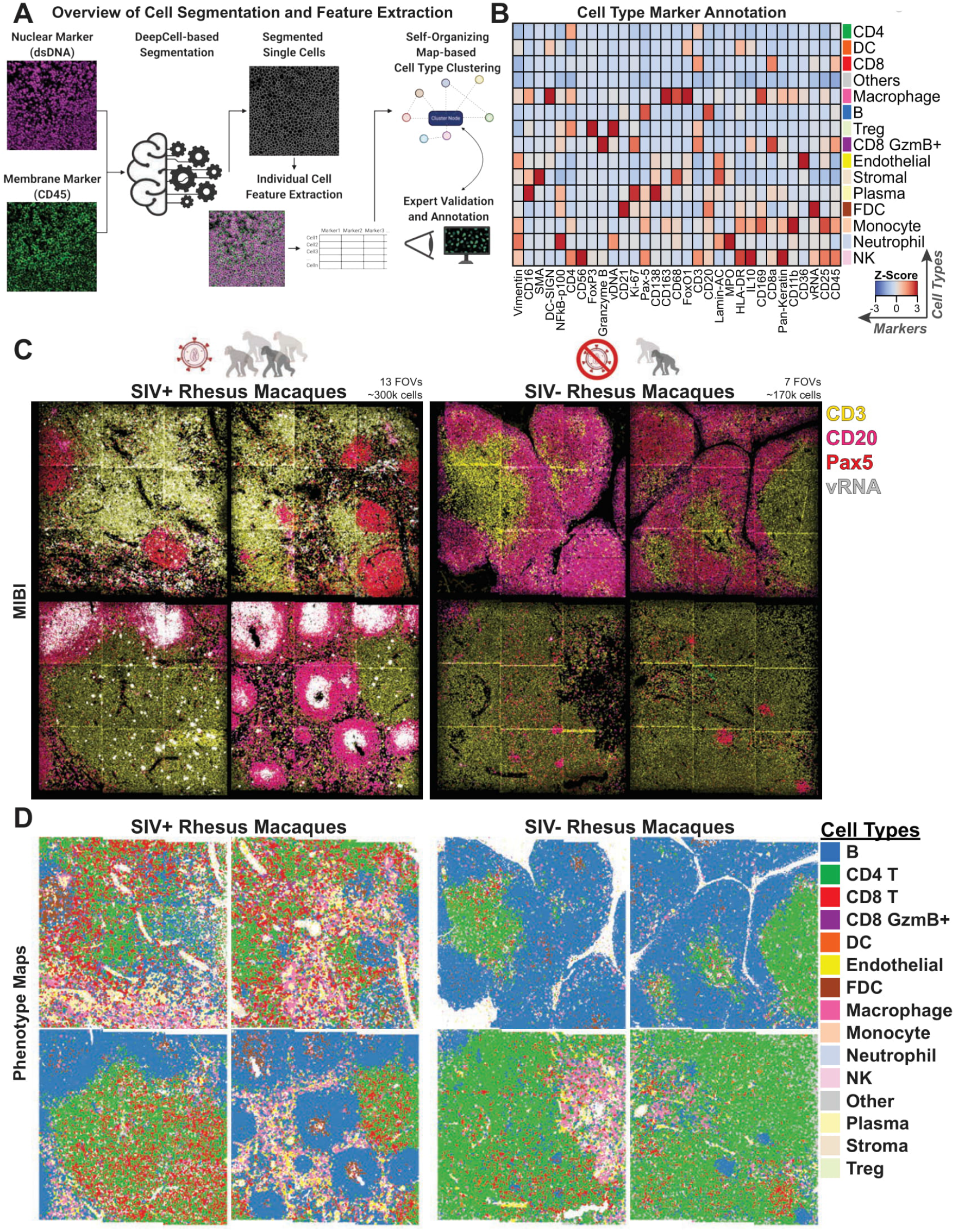
Unsupervised Computation Methods for Feature Extraction and Phenotypic Identification. **(A)** An overview of the deep learning-enabled segmentation of single cells, spatially resolved feature extraction, and self-organizing map-based cell type clustering and annotation used in this study. **(B)** A heatmap depicting the z-scores of marker expression and cell types identified in all FOVs. **(C)** Representative FOVs of tissues from SIV-infected and control animals pseudo-colored to show regions enriched in B and T cells and in SIV vRNA. Each FOV is 1.2 mm x 1.2 mm. A total of 20 FOVs were acquired across four SIV-infected and two uninfected rhesus macaques to generate --470,000 spatially resolved cells. **(D)** Individual cells from the representative FOVs in (C) colored by their cellular phenotypes.

We next analyzed the summary statistics of the 14 different cell types among the 20 individual FOVs (Figure 3A), among the six animals (Figure 3B) and based on the infection status (Figure 3C). There was an evident depletion of CD4^+^ T cells in SIV-infected animals (Figure 3D), a hallmark of HIV-1 and SIV infection (Estes et al., 2008; Picker, 2006; Zeng et al., 2011). B cell numbers were relatively stable, but the infiltration of other immune cell types, such as NK cells, CD8^+^ T cells, FDCs, and macrophages were increased upon infection (Figure 3D). On an individual FOV basis, the amounts of CD8+ T cell, NK cell, and macrophage infiltrations were highly correlated with the infection status of the animal, indicative of the host immune response (Figure 3E). This was also observed for CD8^+^ Granzyme B^+^ T cells, dendritic cells, endothelial cells, FDCs, monocytes, neutrophils, and plasma cells (Figure S4). Extensive SIV deposition was seen on FDCs within B cell follicles in FOVs from SIV-infected tissues, reflective of an expansion of FDCs during infection (Figures 1E, S1C, 2C, and S4). These responses suggest high-order coordination between cell types, beyond phenotypic measurements of individual cells.

**Figure 3:**
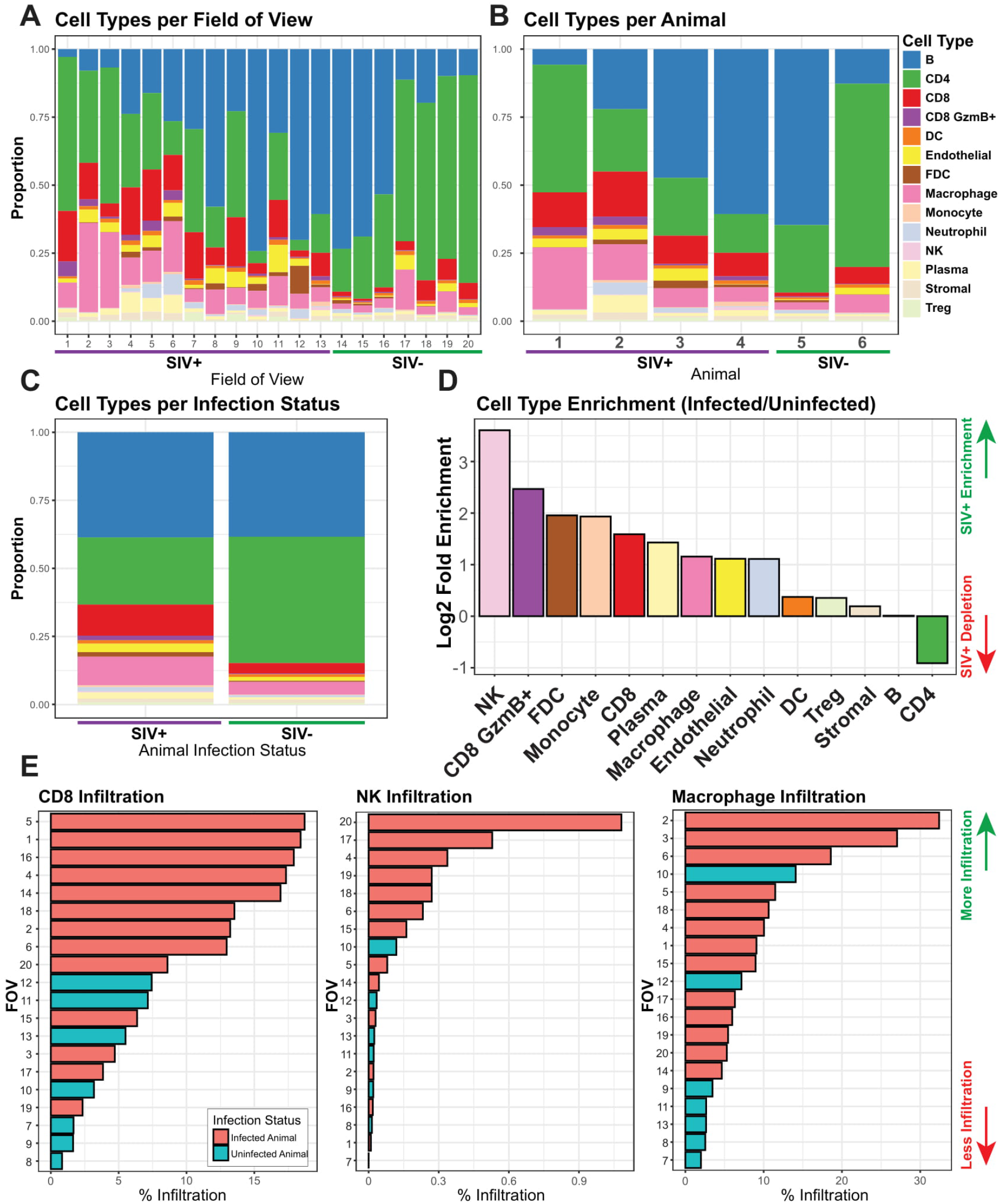
Orchestrated Immune Composition and Responses to SIV Infection. **(A)** Bar plots of proportions of each cell type per FOV across the 20 FOVs acquired in this study. **(B)** Bar plots of proportions of each cell type aggregated on a per animal basis. **(C)** Bar plots representing the proportions of each cell type aggregated by infection status. **(D)** Ranked log2 fold enrichment (infected over uninfected controls) for each cell type, ranked from the most enriched (left) to most depleted (right) in SIV-infected animals relative to uninfected controls. **(E)** Ranked bar plots showing the percent infiltration of each cell type indicated across the 20 FOVs with bars colored by infection status.

### Cellular Neighborhoods Reflect Changes in Tissue Microenvironments upon Viral Infection

Tissue microenvironments are dynamic amalgamations of multiple cell types with ranges of functions within an organ system. Microenvironments are governed by local tissue context such as the immune cell and pathogen composition. Unlike tissue morphologies, which are structural determinants of tissue architecture and the associated cell types (Xu et al., 2009), the tissue microenvironment can be thought of as an accumulation of various chemical and biological determinants exerted both by and onto a cell in its native context. Tissue microenvironments are often qualitatively described by identification of cell types and features, such as blood vessels or immune cells, around a cell of interest. Here we adopted a more empirical Cellular Neighborhood (CN) methodology to quantitatively define the lymphoid microenvironments of healthy and SIV-infected tissues (Schürch et al., 2020). To identify CNs, the cellular phenotypes of the nearest 19 cells around each anchor cell (i.e., 20 cells in total) were quantified and unsupervised clustering was performed. We selected the 20-cell radius as a rough approximation of three cell distances from the anchor cell in each direction, which we visually determined to be a good indication of local functional activity. Thus, CNs take into consideration the impact of the cellular identity of surrounding neighbors on the function of the anchor cell (Figure 4A). Importantly, the infection status of cells and phenotypic and functional markers (e.g., CD4, Ki-67, CD169, and FoxO1) were not considered in defining CNs; therefore, the microenvironment was defined using only the spatial phenotypic patterns. Using this approach, we identified 11 distinct CNs with unique cell compositions (Figure 4B): T cell-, dendritic cell-, and NK cell-rich CN1, B cell zone-containing CN2, macrophage-rich CN3, T cell zone-containing CN4, B cell-, NK-cell, and monocyte-rich CN5, CD4^+^ T cell-rich CN6, FDC-rich CN7, macrophage-rich CN8, stromal and endothelial enriched CN9, CD8^+^ T cell infiltrate-containing CN10, and immune infiltrate-containing CN11. CN summary statistics (Figure 4C) reflect similar trends as the phenotype summary statistics (Figure 3A-D), albeit with additional stratification of cell types such as the CD4^+^ T cells. The ranking of CNs for each FOV revealed the enrichment of certain CNs, including as CNs 5, 7, 10, and 11, in SIV-infected animals (Figures 4D and S5A) and the depletion of the CD4^+^ T cell-rich CN6 (Figure 4C and D).

**Figure 4:**
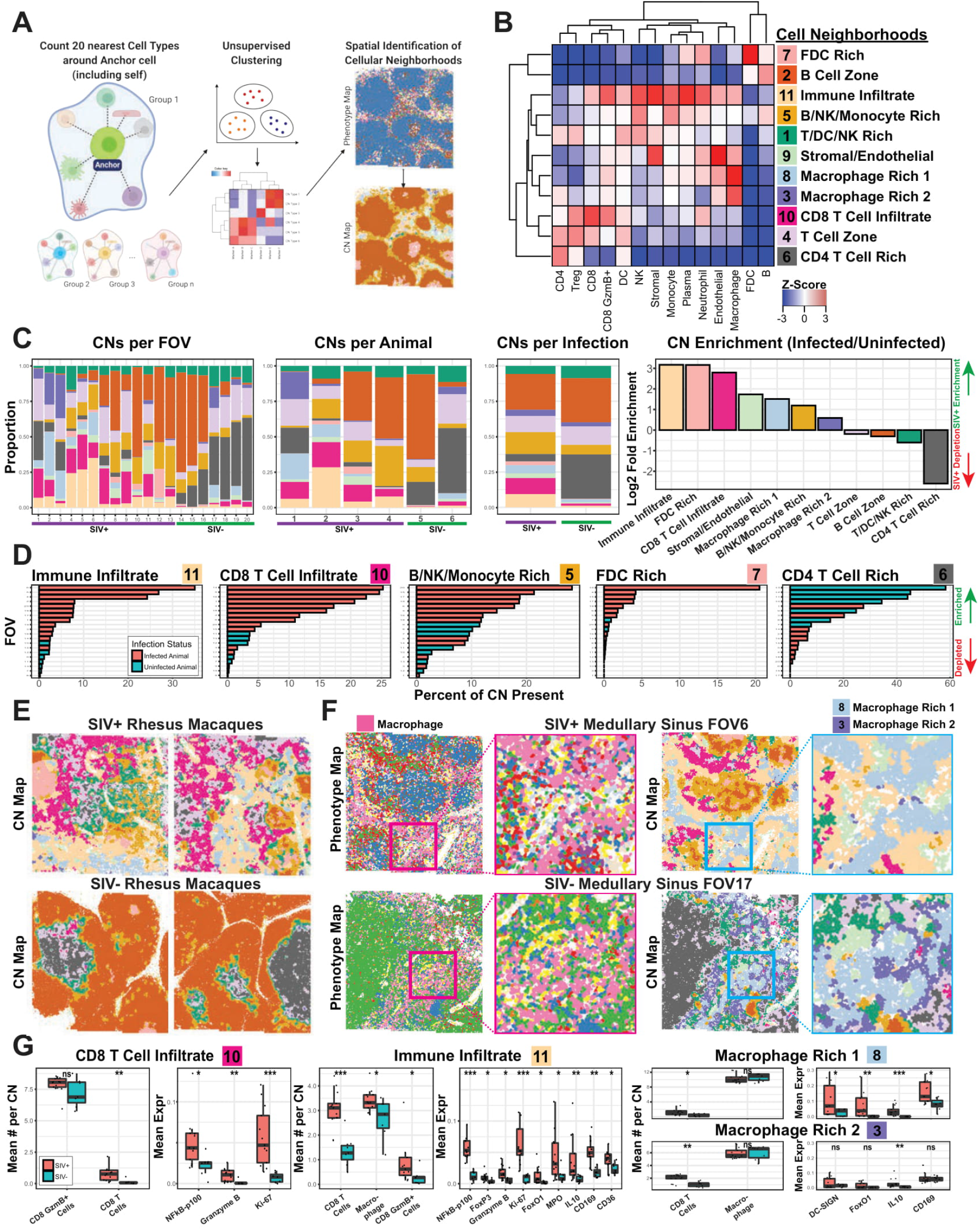
Cellular Neighborhood Analysis Enables Functional Stratification of Tissue Microenvironments during Viral Infection. **(A)** Overview of the method used to define CNs. The 20 nearest neighboring cells (including itself) around each cell were defined, and the cell types identified, quantified, and subjected to unsupervised clustering to define CNs. **(B)** A heatmap depicting the 11 CNs identified and cell types enriched in each, as represented by the z-scores. **(C)** From left to right, bar plots of proportions of each CN aggregated by FOV, animal, and infection status and plot of ranked log2 fold enrichment (infected over uninfected controls) for each CN. **(D)** Ranked bar plots showing the percent composition of each CN across the 20 FOVs with bars colored by infection status. **(E)** Representative FOVs of infected and control animals (also shown in Figure 2C and D) with each individual cell colored by CNs. Each FOV is 1.2 mm x 1.2 mm. **(F)** Representative FOVs from SIV infected (top) and uninfected (bottom) animals containing medullary sinus regions depicted as phenotype maps (left) and CN maps (right). Red and blue boxes indicate regions magnified for zoomed-in views of macrophage-enriched regions. Pink cells in the phenotype map are macrophages. Light blue and purple CNs 8 and 3, respectively, are macrophage-rich. **(G)** Box plots of mean numbers of indicated cell types within CNs (left) and the mean expression of selected functional markers within the CN (right). Each dot in the box plot represents data from a single FOV, and the data are divided between infected (orange) and healthy controls (teal). Non-paired Wilcoxon test: ns, not significant; *, p<0.05; **, p<0.01; ***, p<0.001.

The CN maps are reflective of tissue properties with an additional dimension of information beyond the phenotype maps (Figure 2D, top row, and S3 for phenotype maps; Figure 4E and S5B for CN maps). For example, whereas macrophages predominate in the phenotype maps in SIV-infected and uninfected FOVs, the CN maps for the same areas show a more complex picture with the presence of two different macrophage-rich CNs, CN3 and CN8 (Figure 4F). CN3 (enriched for macrophages and CD4^+^ T cells); is more dominant in SIV-negative FOVs, whereas CN8 (enriched for macrophages, neutrophils, and CD8^+^ GZMB^+^ T cells) is the predominant CN in the SIV-positive FOVs. This highlights that cellular functions are influenced by surrounding external factors.

Quantification of the per cell average levels of SIV vRNA for each CN showed that FDC-rich CN7 had the highest quantities of vRNA (Figure S6), consistent with the major role of FDCs in immune surveillance and antigen presentation. Individual CNs also differed between SIV-infected and uninfected conditions. For example, in FOVs from SIV-infected tissue, for the CD8^+^ T cell infiltrate-heavy CN10, there were more CD8^+^ T cells on average than detected in CN10 regions from healthy control FOVs. We also observed increased CD8^+^ T activation markers such as NFkB-p100 Granzyme B, and Ki-67 (Figure 4G, left). In CN11, which is characterized by the immune infiltrate, both immune cell (CD8^+^ T cells and macrophages) and functional marker levels (NFkB-p100, FoxP3, Granzyme B, Ki-67, FoxO1, MPO, IL10, CD169, and CD36) were elevated in SIV-infected samples compared to uninfected controls (Figure 4G, middle).

Although there were no significant differences between the proportions of macrophages between SIV-infected and uninfected animals in the two macrophage-associated CNs (CN8 and CN3 CD8^+^ T cell abundance was slightly higher in SIV-infected tissue samples (Figure 4G, right). We also observed the dominant expression of macrophage-functional markers (DC-SIGN, FoxO1, IL10, and CD169) specifically in SIV-infected samples in CN8, but only slightly higher expression of IL10 in CN3 in the infected tissue. This suggests that SIV infection induces tissue-specific responses in macrophages. The increase in DC-SIGN, FoxO1, and IL10 (Figure 4G, top tight), markers associated with M2 macrophage antiinflammatory functions, reflects immune dysregulation that occurs during chronic viral infection. The presence of CD169 is reflective of foreign antigen capture by macrophages for presentation. Thus, both phenotypic composition and functional marker expression patterns are altered during viral infection due to immune dysregulation.

### Tissue Architecture Remodeling During Viral Infection

We postulated that CNs can recapitulate the underlying tissue biology as represented by both its cell type composition and functional marker quantifications. We performed linear discriminant analysis (LDA) on the accumulated marker compositions within each CN from each animal (6 animals with 11 CNs from each). LDA analysis separated CNs from infected and uninfected rhesus macaques (Figure 5A, left) and further stratified the animals by chronic versus acute viral infection status (Figure 5A, right). LD1, which accounted for 54.5% of the variation, separated infected and uninfected animals and their associated CNs (Figure 5A and Figure 5B, top). LD2, which captured 22.3% of the variation, distinguished between chronic and non-chronic infection status (Figure 5A and Figure 5B, bottom). Factors differentiating SIV infected from uninfected animals included CD56, CD16, FoxP3, CD11b, and CD36. Factors differentiating SIV chronic from non-chronic status included CD169, CD36, CD16, FoxP3, CD11b, MPO, CD4, CD8a, and Granzyme B. We postulated that the co-occurrences of markers in individual cells would be a proxy to understanding the global tissue reorganization triggered by viral infection. To test this, we calculated the Pearson’s correlations between marker pairs for SIV-negative (Figure 5C, top; teal) and SIV-positive (Figure 5C, bottom; orange) conditions. We focused on markers of cell types dysregulated during SIV infection such as those that characterize B cells, T cells, and macrophages. In agreement with previous data, our strategy highlighted infection-driven processes 1) macrophage immunosuppression as indicated by a M2 switch via CD163 and FoxO1 (blue boxes), 2) increased CD8 T cell infiltration (black boxes), 3) B and T cell proliferation via elevated Ki-67 correlation (yellow boxes) and 4) FDC activation and antigen presentation via increased CD169 and CD11b presence (green boxes).

**Figure 5:**
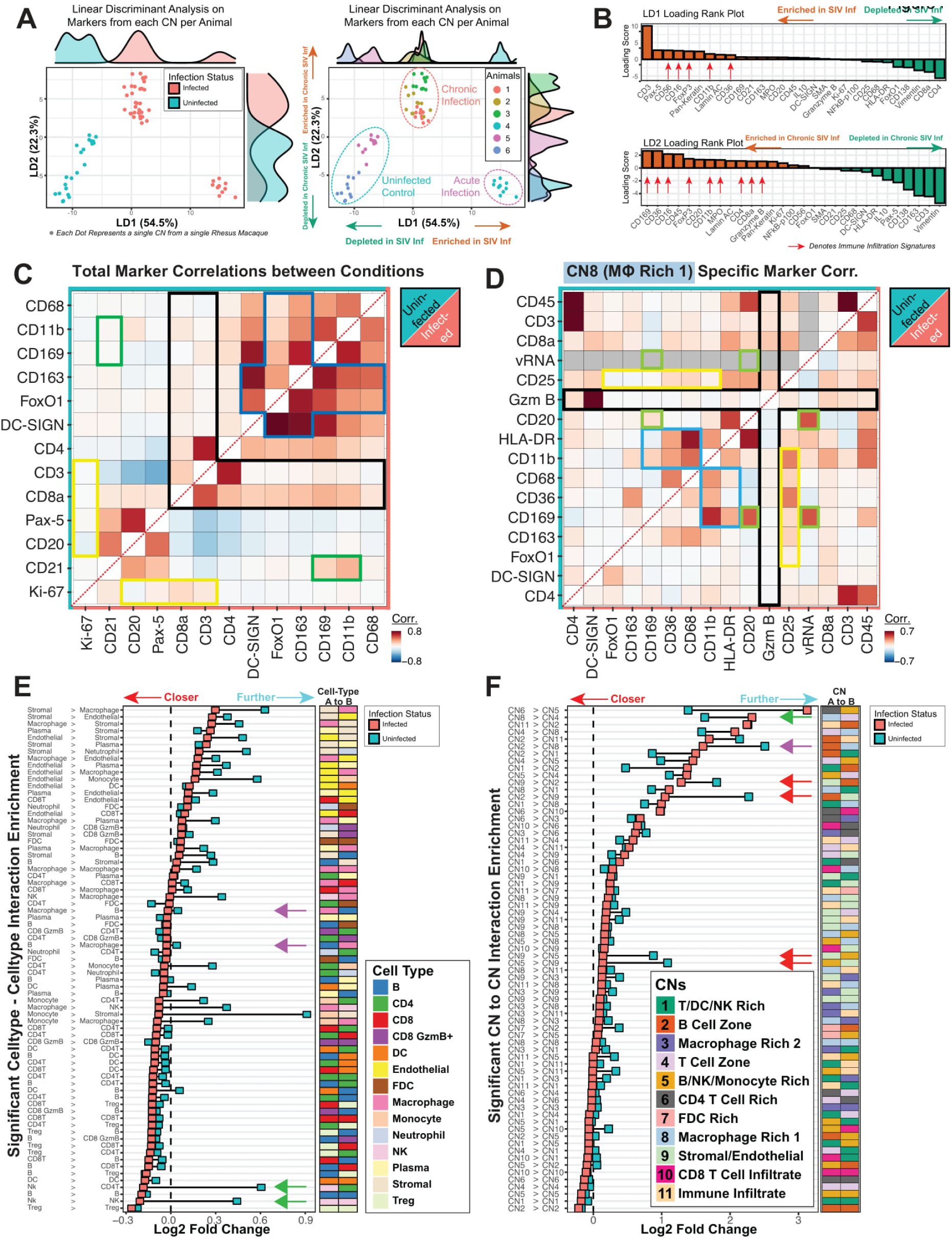
Global and Local Cellular Responses and Tissue Reorganization. **(A)** Plot of variance accounted for by each of the two linear discriminants, LD1 and LD2, on the x and y axes, respectively, for LDA was performed on the collective markers for each of the 14 CNs within each of the six animals. Each dot represents a single CN from a single animal. CNs are colored by the animal infection status (left) or the animal of origin (right). **(B)** The rank plot of the markers that account for LD1 (top) and LD2 (bottom) colored based on enrichment (orange) or depletion (green) in individual CNs of infected animals versus uninfected animals. **(C)** Pairwise Pearson’s correlations of selected immune markers across each individual cell from healthy (top; teal) and SIV-infected (bottom; orange) animals. **(D)** Pairwise Pearson’s correlations of selected immune markers across each individual cell within CN8, separated by healthy (top; teal) and SIV-infected (bottom; orange) animals. **(E and F)** The pairwise cell distances for E) each cell type and F) each CN over randomized background plotted as squares for infected (orange) and healthy (teal) animals. Only interactions that passed a statistical test (p<0.05) for both infection conditions are shown. Squares that are toward the left indicate interactions that are closer than expected, and those toward the right indicate interactions that are further apart than expected. Pairs of cells are given in text form (left) and colored heat maps (right). In panel E, purple arrows indicate B cell-macrophage and macrophage-B cell interactions that are closer in infected versus uninfected tissues, and green arrows indicate NK cell-T cell and T cell-NK cell interactions that are closer in infected than uninfected tissues. In panel F, the green arrow indicates CN8-CN4 interactions that are closer in uninfected than infected tissues, while purple and red arrows indicate CN2-CD8, CN9-CN2, CN2-CN9, CN9-CN5, and CN5-CN9 interactions that are closer in infected tissues than uninfected tissues.

Specific microenvironment interactions were also apparent when Pearson’s correlations between the marker pairs within each CN were analyzed (Figure S7). Notably, for the macrophage-rich CN8 within SIV-positive tissues, there was evidence of 1) increased antigen binding via CD169 but decreased presentation via HLA-DR (blue boxes), 2) decreased granzyme B activity (black boxes), 3) increased CD25 correlation (yellow boxes) and 4) elevated B cell association with vRNA and antigen binding via CD169 (green boxes) (Figure 5D). The pairwise marker correlation maps from single cells in each infection condition and CN provide an informed view of dysregulation and reorganization induced in response to viral infection (Figures 5C, 5D, and S7).

To understand how specific cell types are positioned with purpose and intent within healthy and infected microenvironments, we compared the direction-specific, cell-cell pairwise interactions for each FOV against a randomized background model (Figure 5E). We first identified tissue interactions that were either significantly enriched (Figure 5E, red arrow pointing left) or depleted (Figure 5E, blue arrow pointing right) over the background in both infected and control tissues. Interaction enrichments were then ranked by the SIV-infected status for visualization purposes. For instance, NK-CD4 T cell and NK-NK cell interactions were more likely in SIV-infected tissues than uninfected controls (Figure 5E, green arrows). Both B cell-macrophage and macrophage-B cell interactions were also slightly increased upon infection compared to control tissues (Figure 5E, purple arrows).

We next investigated how CNs were modulated upon viral infection using a direction-specific CN-CN pairwise interaction enrichment analysis over a random background model (Figure 5F). Interactions involving blood vessel-enriched CN9 with CN2 (B cell zone) and with CN5 (B cell-, NK cell-, and monocyte-rich) were prominent in infected tissues (Figure 5F, red arrows), demonstrating increased endothelial interactions and physical proximity of immune cells to specific microenvironments within infected tissues. We also observed that interactions between macrophage-enriched CN8 with CN4 (T cell zone) were decreased in infected tissues (Figure 5F, green arrow), whereas interactions between B cell zone-associated CN2 with macrophage-rich CN8 were increased in infected tissues (Figure 5F, purple arrow). The latter observation is in agreement with the observed increase above in B cell-macrophage and macrophage-B cell interactions (Figure 5E, purple arrows). It is important to note that although CN2 and CN8 were closer in SIV-infected tissues than uninfected tissues, these neighborhoods did not physically interact or overlap (Figure S5B).

### IL10-Induced Immunosuppressive Microenvironments by B Cells and Macrophages

Our results suggest that viral infection induces a strong linkage between B cells and macrophages (Figure 5E, purple arrows), specifically between CN2 (B Cell Zone) and CN8 (Macrophage Rich 1) (Figure 5F, purple arrow). The IL10 expression patterns were distinctive in these neighborhoods: cells positive for IL10 were predominantly B cells in CN2 (Figure 6A, left) and were predominantly macrophages in CN8 (Figure 6A, right). IL10 is an immunoregulatory cytokine that can activate or suppress the immune system (Ouyang and O’Garra, 2019; Rojas et al., 2017). IL10 expression is upregulated in HIV patients within several circulating immune cell types, including B cells (Brockman et al., 2009), and in lymphoid tissues after SIV infection (Estes et al., 2006; Tabb et al., 2013). Both B cells in CN2 and macrophages in CN8 had increased IL10 expression after SIV infection (Figure 6B), implicating elevated IL10 expression by B cells and macrophages as a host response to viral infection. We observed a positive correlation between vRNA and IL10 levels within CN2 (Figure 6C, top left panel) but no correlation within CN8 (Figure 6C, bottom left panel). These results suggest that either extensive deposition of viral particles on FDCs, viral progeny production from infected cells, or both are upstream of IL10 production in the B cell-rich CN2 and are downstream of IL10 production in the macrophage-rich CN8.

**Figure 6:**
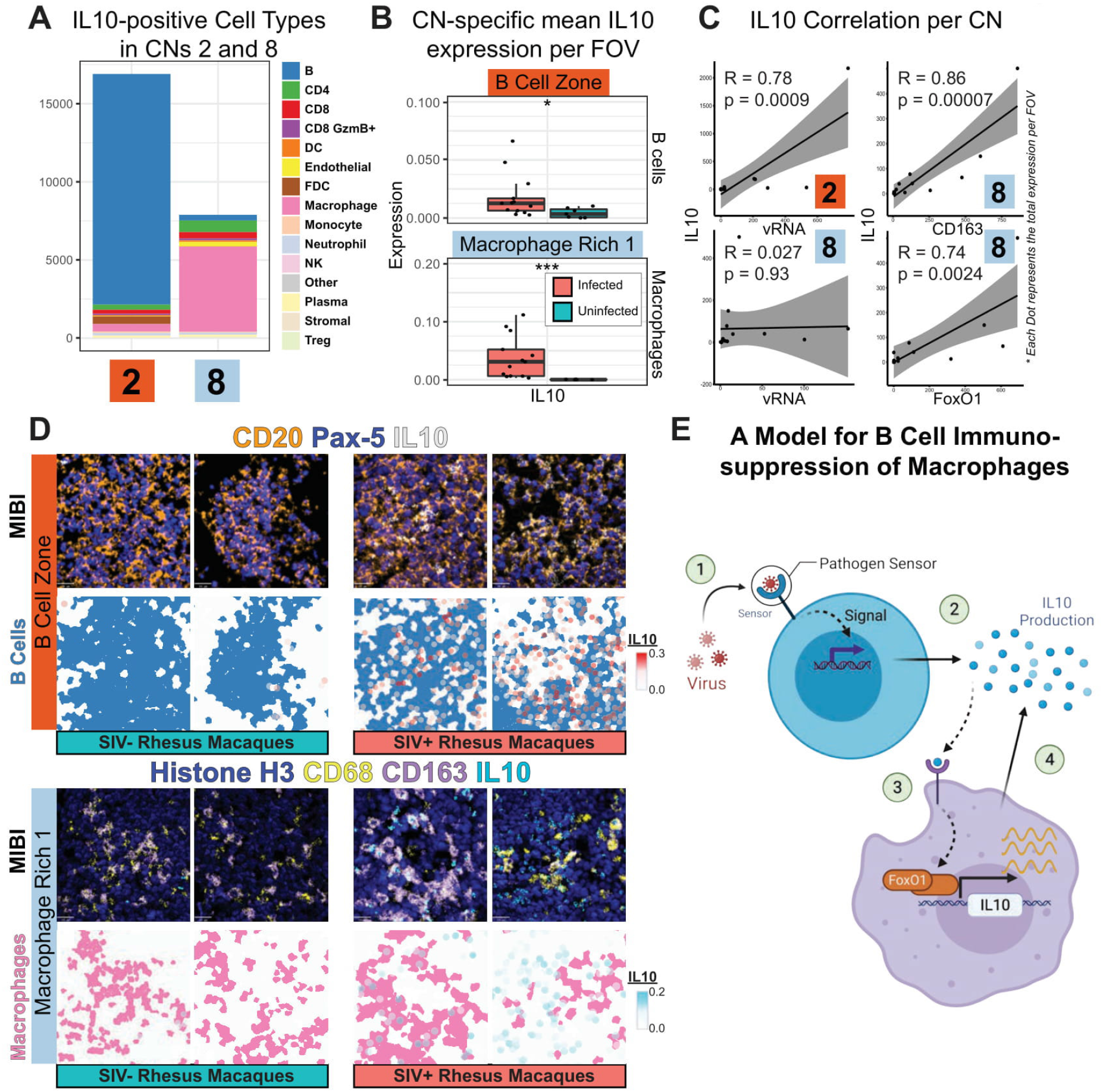
B Cell-Driven IL10 Production Results in Immunosuppression during SIV Infection. **(A)** Bar plot of numbers of IL10-positive cells of the indicated types in CN2 and CN8 in all animals. **(B)** Box plots of mean IL10 expression across B cells within the CN2 and across macrophages in CN8. Each dot represents data from a single FOV from SIV-infected and uninfected controls. **(C)** Plots of Pearson’s correlations between IL10 levels and vRNA in CN2 (top left), vRNA in CN8 (bottom left), M2 immunosuppressive macrophage marker CD163 in CN8 (top right), and FoxO1 in CN8 (bottom right). Each dot represents data from a single SIV-infected FOV. **(D)** Representative pseudo-colored MIBI images depicting IL10 and B cell markers CD20 and Pax-5 and IL10 (top) and IL10 and macrophage markers CD68 and CD163 (bottom). Two representative FOVs from infected and uninfected animals are shown. The phenotype maps superimposed with IL10 expression patterns are shown below each MIBI image. **(E)** A cartoon depicting the model for B cell-induced immunosuppression of macrophages via IL10. 1) B cells sense SIV virions via an unknown receptor. 2) B cells produce IL10, and possibly other chemokines to attract nearby macrophages. 3) Binding of IL10 to macrophages activates downstream factors, including FoxO1, which leads to more IL10 production and release. 4) This feed-forward loop results in an immunosuppressive microenvironment around these macrophages.

Given the ability of IL10 to suppress immune cells, we next measured the relationship between levels of IL10 and immunosuppressive M2 macrophage markers CD163 and FoxO1 in CN8. There was a positive correlation between these markers (Figure 6C, right panels), implicating IL10 in triggering of M2 macrophage differentiation. We visually confirmed our findings pertaining to the upregulation of IL10 within B cells (Figure 6D, top) and CD163^+^ M2 macrophages (Figure 6D, bottom) in response to SIV infection. Together, these results support a model for how SIV infection purposefully induces immunosuppressive TMEs and cell phenotypes through 1) initial sensing of viral particles by B cells (perhaps through CD21/CR2 engagement), 2) production of IL10 that then attracts nearby macrophages, 3) subsequent FoxO1 activation that leads to more IL10 production and finally 4) M2 macrophage differentiation and creation of an immunosuppressive TME (Figure 6E).

### Environmental Cues influence SIV Viral Latency

PANINI enables detection of vDNA and vRNA, with host proteins, and is therefore particularly suitable for the detection of SIV-infected cells that are latent (i.e., vDNA^+^ and vRNA^−^) and those that are transcriptionally active (vDNA^+^, vRNA^+^). We identified 914 SIV-infected cells within rhesus macaque lymph nodes. These cells were predominantly CD4^+^ T cells (69.7%), of which 10.3% were Tregs and the rest macrophages (30.3%) (Figure 7A). Consistent with the cohort of viremic SIV acute and chronically infected animals, the infected cells were predominately transcriptionally active (64.4%), with a similar composition of latency status within each cell type and CN of origin (Figure 7A). The only exception was the higher presence of latent cells within CN9, which was enriched in stromal and endothelial cells (Figure 7A).

**Figure 7:**
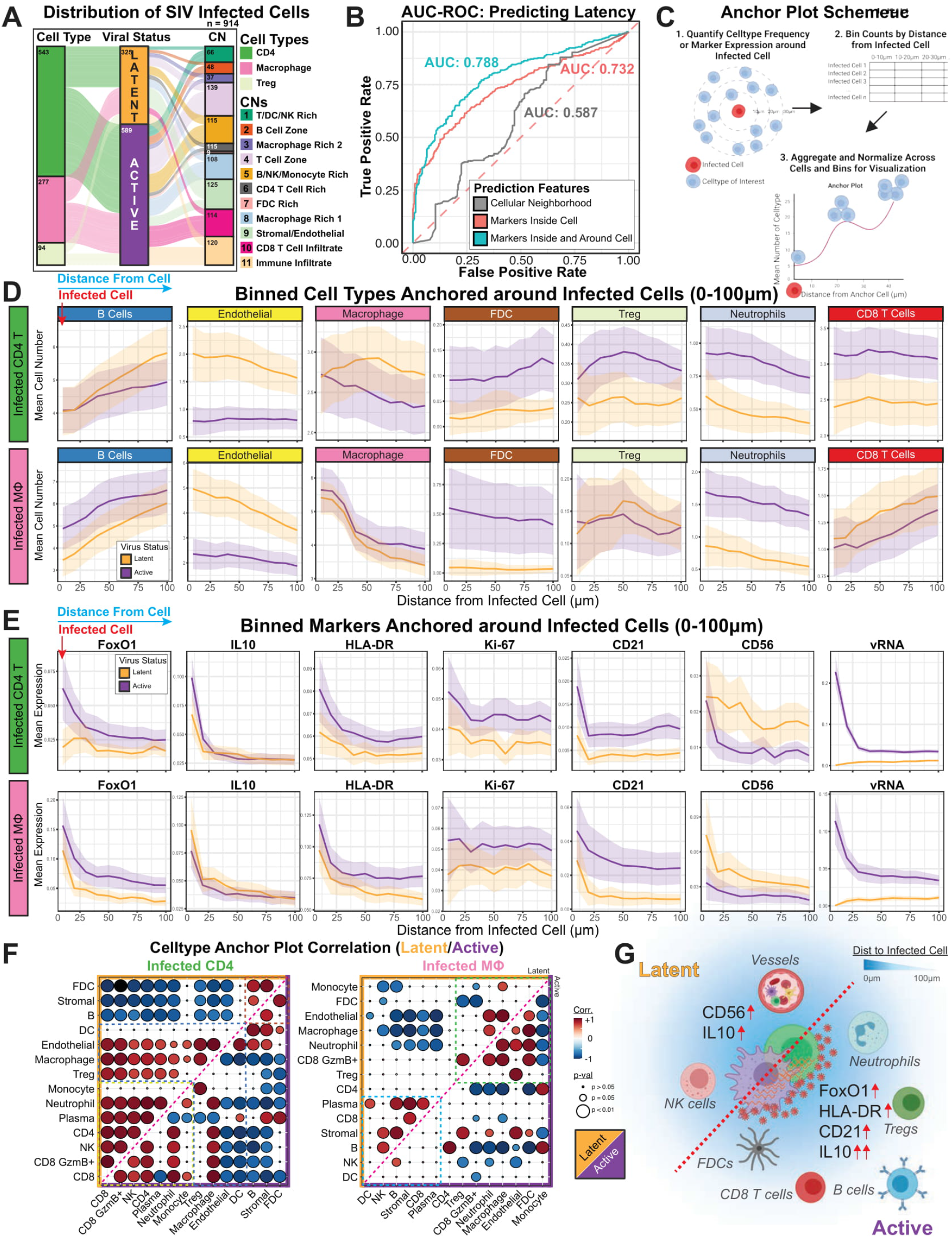
Host Determinants of Retroviral Latency. **(A)** An alluvial plot depicting the compositions of SIV-infected cell types (CD4^+^ T cells, macrophages, and Tregs), their latency status, and their associated CNs. **(B)** Predictive performances of classifiers 1) CNs (gray, AUC = 0.587), 2) markers inside the infected cell (orange, AUC = 0.732), and 3) markers from a cell and its nearest neighbors (teal, AUC = 0.788). The dotted red line indicates AUC of 0.5, the expected by chance. **(C)** A schematic depicting how the anchor plots were calculated for the anchor plots in panels D and E. In short, 1) mean cell type frequencies or marker expressions around each infected cell were tabulated, 2) these values were binned by their distance from the infected cell in 10-μm increments, and 3) data for all infected cells were aggregated and normalized for visualization. **(D and E)** Anchor plots of D) mean cell type quantifications and E) mean marker expression around infected CD4^+^ T cells (top) or macrophages (bottom). Orange indicates latent cells, and purple indicates actively transcribing cells. The thick colored lines represent the means, and light regions around these lines depict the 95% confidence intervals. The infected cells are anchored at 0 μm, and the plot ends at 100 μm. **(F)** Heatmaps of Pearson’s correlations for cell type pairs for infected CD4^+^ T cells (left) and macrophages (right). Latent infection correlation heatmaps are represented to the top left (orange)and active infection correlation heatmaps are in the bottom right (purple). The sizes of the circles reflect the p-values from the test for association using the correlation coefficient, and colors indicate degree of correlation (legend on the right). **(G)** Schematic representing the tissue correlates and determinants of retroviral latency status in retroviral reservoirs.

To identify cellular and CN features predictive of viral latency, we trained a random forest classifier on 1) CN information alone, 2) cell marker features within the infected cell, and 3) cell marker features within the 20 cell-radius (i.e., 19 neighbors and the infected cell). Viral RNA and NFkB-p100 were excluded from the prediction features as the former are directly related to the definition of latency status while the latter is critical for vRNA transcription (Hiscott et al., 2001). We observed that CNs alone were poor predictors of viral latency, with the area under the curve (AUC) of the receiver operating characteristics curve of 0.587, close to what is expected by chance (Figure 7B). Markers within the infected cell were better predictors of latency than CNs (AUC: 0.732), whereas utilizing markers of the infected cell and its neighbors resulted in the best predictive performance (AUC: 0.788). These observations indicate that factors both intrinsic to the infected cell and those from the environment influence viral latency status.

How cells communicate through cell-to-cell interactions and soluble mediators are products of their proximity to each other and the marker expression patterns in their vicinity. We devised a meta-analytical method to quantify both the celltype frequency and marker expression around centered cells of interest. In short, we quantified cell types and their marker expressions within a 100-μm radius around aggregated infected cells and then split them into ten bins for normalization, plotting, and visualization (Figure 7C). We focused on the more abundant non-Treg infected CD4^+^ T cells and macrophages, given the low number of infected Tregs (n = 94) (Figure 7A). We observed that infected cells tended to be within regions with high B cell density regardless of latency status (Figure 7D). Furthermore, latent cells were generally much closer to endothelial cells than were transcriptionally active infected cells (Figure 7D), indicative of how a constant influx of naïve immune cells into the tissue reinforced a viral latent state for immune evasion. Infected cells were also in proximity to macrophages, although this did not depend on viral latency status or type of infected cell (Figure 7D). Interestingly, there were higher densities of FDCs, Tregs, neutrophils, and CD8^+^ T cells in the vicinities of infected CD4^+^ T cells actively producing vRNA than in the vicinities of latent CD4^+^ T cells (Figure 7D). This was also observed for transcriptionally active infected macrophages with the exception of proximal Tregs and CD8^+^ T cells (Figure 7D). The higher densities of FDCs around infected cells actively transcribing viral RNA is reminiscent of experiments in which isolated FDCs were capable of dramatically augmenting HIV transcription *in vitro* (Thacker et al., 2009).

Quantifying functional markers enables insights into an additional layer of microenvironmental complexity beyond mere cell phenotypes. vRNA expression patterns around cells actively transcribing vRNA followed a clear diffusion pattern from the source (Figure 7E, right), and we observed a stark scaling of the amount of FoxO1 expression as a function of distance from active versus latent cells (Figure 7E). IL10 levels tended to be elevated around all infected cells and were even higher around active versus latent CD4^+^ T cells (Figure 7E). Similar trends were observed for HLA-DR, Ki-67, and CD21 where higher proximal expressions correlated with viral activation within infected cells (Figure 7E). The reverse was observed for CD56, where elevated levels around latently infected cells suggested of a role of pathogen recognition by CD56 expressed on NK cells in controlling viral latency (Figure 7E). Surprisingly, we detected similar and distinctive patterns for various markers in infected CD4^+^ T cells and macrophages (Figure 7D-E and S8), emphasizing the importance of recognizing both the cellular phenotype and functional markers around infected cells and their microenvironment (Figures 7D, 7E and S8).

### Orchestrated Events Condition Tissue Microenvironments During Infection

To better interpret the complexity of orchestrated tissue events that may determine the viral activation status of infected cells, we computed the Pearson’s correlations between each pair of binned cell type frequencies as a function of distance from the infected cell (Figure 7F; top, yellow: latent cells, bottom, purple: active cells). Three modules were detected in infected CD4^+^ T cells (Figure 7F, left). One module of interaction involving FDCs, stromal cells, and B cells, factors essential for germinal center functions, was disrupted during viral RNA transcription in infected CD4^+^ T cells (Figure 7F, left). Another module populated by dendritic cells, endothelial cells, macrophages, and Tregs was anti-correlated in active infected CD4^+^ T cells (Figure 7F, left). The third module composed of monocytes, neutrophils, plasma, CD4^+^ T cells, NK, and CD8^+^ T cells also differentiated CD4^+^ T cell latency status (Figure 7F, left). For infected macrophages, we observed two signature modules (Figure 7F, right). The first was dominated by monocytes, FDCs, macrophages, neutrophils, endothelial, CD8^+^ Granzyme B^+^ cells and Tregs. The second involved predominantly plasma cells, CD8^+^ T cells, stromal cells, B cells, NK cells, and dendritic cells. Taken together, these plots show that the fundamental cellular relationships in the vicinity of infected, latent cells are disrupted when compared to infected cells actively producing viral RNA. The interplay between virus-infected cells and the proximity of microenvironmental signals reveal roles for both intrinsic and external factors in shaping the tissue microenvironment as effectors or consequences of viral infections. Our data support a model in which the distances of both functional markers (e.g., CD56, IL10, FoxO1, HLA-DR, and CD21) and cell types (e.g., endothelial vessels, NK, FDCs, neutrophils, Tregs, B cells, and CD8^+^ T cells) from SIV-infected cells have a strong influence on latency status (Figure 7G).

## Conclusions

Here we described development and validation of PANINI, a robust platform that enables detection of nucleic acids with high sensitivity while preserving confident detection of protein epitopes. This was achieved through a combination of heat-induced epitope retrieval in a pH9 buffer, branched-chain amplification of nucleic acids without the need for protease treatment, subsequent tyramide signal amplification coupled with hapten deposition, followed by multiplexed antibody staining for protein and nucleic acid (via anti-hapten Ab) recognition. In this study, we coupled PANINI with the MIBI to spatially detect integrated viral DNA, viral RNA that is present in actively replicating cells and viral particles (vRNA), along with 31 immune phenotypic and functional protein markers (Figure S1). We demonstrate the utility of PANINI in the detection of nucleic acid copies down to single events in archival FFPE tissues, which has been notoriously difficult to study when coupled with protein detection methodologies. This uniquely enabled interrogation of the diverse immune responses within SIV-infected lymphoid tissues, particularly the uncharted phenotypes and spatial features around latently infected cells and around cells in which viral RNA is being transcribed.

We first confirm hallmarks of retroviral infection, such as the depletion of the CD4 T cell population (Figures 3C and 3D) (Hazenberg et al., 2000), a heightened NK cell and CD8 T cell response (Figure 3E) (Alter et al., 2007; Goonetilleke et al., 2009), and a lack of immune infiltration into the “sanctuary” B cell follicles (Figures 4B and 4E) (Fukazawa et al., 2015). Previous studies have also highlighted the upregulation of IL10, a powerful cytokine capable of dampening an immune response, in multiple immune cells within PBMCs of HIV-infected individuals, though its role and distribution in tissues is largely unclear (Brockman et al., 2009; Estes et al., 2006; Fukazawa et al., 2015; Tabb et al., 2013). Here, leveraging upon the capability of PANINI to robust nucleic acid detection, with the highly multiplexed imaging capabilities of the MIBI, we reveal a B cell response to SIV-infection through the secretion of IL10, in addition to likely other cytokines. This is followed by the attraction and immunosuppression of macrophages in its vicinity via infection of a M2-phenotypic switch, thus organizing an immunosuppressive microenvironment in SIV-infected tissues (Figure 6). Such a dampened environment appears to harbor viral-infected cells, with heightened IL10, FoxO1 and HLA-DR expression ~20-30um around the infected cell stratifying in part its latency status (Figure 7). This is just one example of how our data highlights the temporal ordering of distinctive tissue features during SIV infection that warrant further investigation in suitable animal models.

Robust multiplexed imaging of nucleic acids, concurrently with proteins, is particularly suited for disentangling environmental effects from intrinsic properties of the cell. We demonstrated this using SIV-infected rhesus macaques as a model, with a particular focus on viral reservoirs within lymphoid tissues, sites previously described to be a primary location of infected cells (Estes et al., 2017). We confirmed the robustness of our assay by identifying CD4^+^ T cells and macrophages as the primary cell types infected and discovered that both extrinsic and intrinsic features are required to predict viral activation status best. For example, using a distance-based analysis anchoring around infected cells, we uncover how infected cells 1) tend to be in regions densely populated with B cells, previously described as immune-privileged “sanctuaries”, 2) tend not actively transcribe viral genes when situated in close proximity to vasculature, and 3) tend to produce viral transcripts when in close proximity to FDCs. Additional contributing features that distinguish between viral latency and transcription include the expression of CD56, and quantities of Tregs, neutrophils and CD8 T cells (Figure 7G).

The establishment of the PANINI experimental platform, a 33-marker panel compatible with FFPE archival tissues, spatial analytical workflow, and conceptual framework for multimodal analysis of tissue features have enabled reinterrogation of previous observations and establishment of new models and hypotheses. However, fundamental questions remain, including: 1) How do CNs and infected cells change with antiretroviral therapy (ART) or immunotherapy? 2) Are features and relationships different in other tissue sites, such as the brain or gut-associated lymphoid tissue? 3) Can these principles be translated to other infectious diseases such as tumor virus-driven malignancies, SARS-CoV-2, and Tuberculosis, or cancer biology questions involving copy-number amplifications, repetitive elements and extra-chromosomal DNA? We anticipate PANINI, coupled with widely adopted multiplexed imaging technologies such as MIBI, CODEX, cycIF, and IMC, well-validated nucleic acid probes and antibodies, and robust animal models or archival clinical samples, will be essential for advancing the mechanistic insights needed to better guide therapeutic intervention strategies.

## Supporting information

Key Resource Table

## ACKNOWLEDGEMENTS

The authors thank previous and current members of lonpath Inc, including MatthewNewgram and Maciej Zerkowski, for their unwavering technical support. We thank members of the Nolan, Estes and Angelo labs for helpful discussions. S.J. is supported by the Leukemia & Lymphoma Society Career Development Program and a Stanford Dean’s Fellowship. X.R.C. is supported by the EMBO postdoctoral fellowship (ALTF 300-2017). B.Z. is supported by a Stanford Graduate Fellowship. This work was funded in part by grants from the National Institutes of Health R01AI149672 (G.P.N. and J.D.E.), the Bill & Melinda Gates Foundation INV-002704 (G.P.N. and J.D.E.), OPP1113682 (G.P.N.), P51OD011092 (J.D.E.) and COVID-19 Pilot Award (S.J., D.R.M., G.P.N.), the Fast Grant Funding for COVID-19 Science (G.P.N.), the US Food and Drug Administration Medical Countermeasures Initiative contracts HHSF223201610018C and 75F40120C00176 (G.P.N.), the Parker Institute for Cancer Immunotherapy (G.P.N.), and the Rachford and Carlota A. Harris Endowed Professorship (G.P.N.). This article reflects the views of the authors and should not be construed as representing the views or policies of the FDA, NIH, BMGF or other institutions who provided funding.

## Data availability

All multiplexed imaging data produced in this paper are available via the MIBItracker Public Accession Portal at the following link: https://mibi-share.ionpath.com/

## AUTHOR CONTRIBUTIONS

Conceptualization: S.J., J.D.E.

Methodology: S.J., J.D.E.

Experimental Investigation: S.J., C.N.C., X.R.C., H.C., Y.H., B.Z., J.P.O., N.M., J.D.E.

Computational Analysis: S.J., H.C., Y.H., C.L.

Novel Reagents and Tools: E.M., N.G., G.L.B., J.L.W., D.P., K B.S., M.N., M.T., S.Y., M.B., J.D., Y.G.

Visualization: S.J., H.C., C.N.C.

Writing: S.J., C.N.C., X.R.C., J.D.E., G.P.N.

Funding Acquisition: S.J., D.R.M., J.D.E., G.P.N.

Supervision: S.J., D.R.M., M.A., J.D.E., G.P.N.

All authors read and approved the final draft of the manuscript.

## Materials & Methods

### Animal Experiments and Tissue Acquisition

Archival FFPE tissues were obtained from SIV-infected and control rhesus macaques (Macaca mulatta) of Indian origin that were housed at the Oregon National Primate Research Center (OR, USA) and at the National Institutes of Health (Bethesda, MD, USA) with the approval of the respective Institutional Animal Care and Use Committees. The animal experiments were conducted with strict adherence to the NIH and the Animal Welfare Act and in accordance with American Association for the Accreditation of Laboratory Animal Care (AAALAC) standards in AAALAC-accredited facilities.

Our cohort consisted of the following animals: SIV-negative (n=2) and SIV-challenged (13 day or 16-19 weeks post infection; n=4). Lymph nodes were collected at necropsy and immediately fixed in freshly prepared neutral buffered 4% PFA for 24 hours at room temperature. Afterwards, the fixative was replaced with 80% ethanol and the tissues were processed through a series of 30-minute incubations in increasing alcohol concentrations to 100 percent, then in xylene and hot paraffin, in a Tissue Tek Vacuum Infiltration Processor 6 (Sakura). Processed tissues were then paraffin embedded and stored in a cool, dry place.

### Antibody Conjugation

Antibodies were conjugated to metal polymers using the Maxpar X8 Multimetal Labeling Kit (Fluidigm, 201300) and Ionpath Conjugation Kits (Ionpath, 600XXX) with slight modifications to manufacturer protocols. The antibodies used and their respective clones are listed in the Key Resources. Antibody conjugation was performed exactly as described previous in Han et al (Han et al., 2018). In short, 100ug of carrier free antibodies are subject to gentle reduction in the presence of 4uM of TCEP for 30 min, before conjugation to lanthanide-loaded polymers. Post elution, all antibodies are quantified via nanodrop (Thermo Fisher Scientific, ND2000), diluted into >30% w/v Candor PBS antibody stabilizer (Thermo Fisher Scientific, nc0436689) and stored at 4°C until use.

### Gold Slide Preparation

Gold slides were prepared as previously described (Ji et al., 2020; Keren et al., 2018). Briefly, Superfrost Plus glass slides (Thermo Fisher Scientific, #12-550-15) were soaked in dish detergent, rinsed with distilled water followed by acetone. Acetone evaporation was performed under a constant stream of air in a fume hood, and clean slides subsequently coated with 30nm of Tantalum followed by 100nm of Gold at the Stanford Nano Shared Facility (SNSF; Stanford CA) and New Wave Thin Films (Newark, CA).

### Vectabond Pre-treatment of Gold Slides

Gold slides were pretreated with Vectabond (Vector Labs, #SP-1800) according to the manufacturer’s protocols. In short, slides were submerged in 100% acetone for 5 min before incubation in a glass beaker containing a mixture of 2.5 ml Vectabond and 125 ml 100% acetone for 30 min. Slides were then washed in 100% acetone for 30 sec, air dried, and stored at room temperature.

### Cell Culture and FFPE Cell Pellet Embedding

The well-characterized SIV-infected cell line 3D8, which contains a single integrated provirus per cell (Mattapallil et al., 2005), and the uninfected parental 174xCEM cell line were used to validate our detection of vRNA and vDNA as part of the PANINI workflow (see below). Cell were fixed in 4% paraformaldehyde (PFA) overnight before embedding into Histogel (Fisher Scientific, NC9150318) and paraffin wax as described previously (Deleage et al., 2016).

### RNAScope & DNAScope Fluorescent Multiplex *in situ* Hybridization

The RNAScope & DNAScope multiplex staining methodology originally described in (Deleage et al., 2016) was modified and optimized to increase the feasibility of using a pH9 antigen retrieval condition without protease digestion to detect both SIV vDNA and vRNA. FFPE sections of SIV-positive and SIV-negative rhesus macaque lymph nodes on Fisher Superfrost glass microscopic slides were deparaffinized by heating at 60°C for 1h and then transferred to a xylene bath for 5 mins. Slides were transferred to a new xylene bath for another 5 min, followed by 2 x 1 min incubations in 100% EtOH baths. Slides were then rinsed with double distilled water (ddH2O) and boiled in 1X Dako pH9 antigen retrieval solution (Agilent, S236784-2) for 10 min. The slides and the hot retrieval solution were left to cool down at room temperature for another 20 min before the slides were rinsed twice in ddH2O. A hydrophobic barrier was drawn around the tissue using the ImmEdge Hydrophobic Barrier pen (Vector Labs, 310018). For slides that were treated with Protease, the tissue was treated with Protease III (Biotechne, 322337) diluted 1:10 with cold PBS and incubated at 40°C in an ACD HybEZ Hybridization System oven (Biotechne, 310013) for 20 min, then rinsed twice with ddH2O. Slides not treated with protease remained in ddH2O throughout this process. Next, endogenous peroxidase was inactivated using 3% H2O2 in PBS and rinsed twice in ddH2O.

Slides were incubated overnight at 40°C with RNAScope probes that detect SIVmac239 vif-env-nef-tar vRNA (Biotechne, 416131-C2) and SIVmac239 gag-pol vDNA (Biotechne, 416141). The next day, slides were washed twice with 0.5X Wash buffer (Biotechne, 310091) for 2 min each. Branched-chain amplification was performed using the Multiplex Fluorescent V2 kit (Biotechne, 323110) with the following conditions, with a 2 x 2 min wash between each step:

- Amplifier 1, 30 min at 40°C
- Amplifier 2, 15 min at 40°C
- Amplifier 3, 30 min at 40°C
- Channel 1 specific:

1. Amplifier 4, 15 min at 40°C
2. Custom Amplifier 5, 30 min at room temperature
3. Custom Amplifier 5, 15 min at room temperature
4. Biotium Tyramide CF640R deposition, 15 min at room temperature
- Hydrogen peroxidase block (Biotechne, 323107), 15 min at 40° C
- Channel 2 specific:

1. Amplifier 4, 15 min at 40°C
2. Custom Amplifier 5, 30 min at room temperature
3. Custom Amplifier 5, 15 min at room temperature
4. Biotium Tyramide CF568 deposition, 15 min at room temperature

All TSA hapten reagents were diluted in an in-house TSA diluent (0.1M Borate, pH 8.5, with 2% w/v Dextran Sulfate Sodium salt and 0.003% H2O2, with the H2O2 added just before the dilution of the tyramide reagent) for 3 minutes at room temperature. We observed that in slides without protease treatment, a higher concentration of CF568 was needed to fully amplify vRNA signals (as determined from FDC-bound SIV vRNA) compared to protease-treated slides. It is important to note that we did not observe any other differences, such as off-target signals and tissue morphological changes, from this increased concentration of CF568. Slides were then rinsed once with 1 x TBS-T, counterstained with DAPI and cover-slipped with #1.5 GOLD SEAL cover glass (EMS, 63791-10) using Prolong Gold Mounting medium (ThermoFisher, P36930). Whole-slide high-resolution fluorescent scans were performed using a Plan-Apochromat 20X objective (NA 0.80) in the Zeiss AxioScan Z.1 slide scanner. DAPI, AF568 and Cy5 (For CF640R) channels were used to acquire images. The exposure time for image acquisition was between 4 and 300 ms.

### PANINI Staining

FFPE tissue paraffin blocks were sectioned onto vectabond treated gold slides at 4 um thickness on a microtome. Slides were baked for 1h at 70°C and soaked in xylene for 3 x 10min. Standard deparaffinization was performed thereafter (3 x xylene, 3 x 100% EtOH, 2 × 95% EtOH, 1 x 80% EtOH, 1 × 70% EtOH, 3 x H2O) on a linear stainer (Leica Biosystems, ST4020). Epitope retrieval was performed at 97C for 10 min with the Dako Target Retrieval Solution pH 9 (Agilent, S236784-2) on a Lab Vision PT Module (Thermo Fisher Scientific).

Slides were cooled down to 65°C in the PT Module, and left to further cool to room temperature. The region containing the tissue sections were traced out using an ImmEdge PAP pen (Vector Labs, H-4000) before rinsing 2 x 2 min in ddH2O. Tissue sections were then subject to a hydrogen peroxidase block (Biotechne, 322330) at 40°C for 15 min, before 2×2 min ddH2O wash. Avidin and Biotin blocks (Biolegend, 927301) were then performed for 15 min each at room temperature, with 2 x 2 min ddH2O washes after each block.

RNAscope probes (see Key Resources) were then added for overnight hybridization (~18 hrs), and all washes from hence forth were performed using RNAscope wash buffer (Biotechne, 310091) for 2 x 2 min at room temperature. Branched-chain amplification using a customized version of the Multiplex Fluorescent Detection Kit v2 (Biotechne, 323110), in which additional amplification was enabled (Amplifiers 5 & 6) for each channel. All amplification reactions were performed at 40°C, with the exception of the following which occur at room temperature: 1) Amplifiers 5 & 6 and 2) Hapten-deposition via tyramine signal amplification (TSA). Reagents for TSA hapten deposition were Biotin (Akoya, NEL749A001KT) and DIG (Akoya, NEL748001KT). All 40°C steps were performed in an ACD HybEZ Hybridization System oven (Biotechne, 310013).

Branched-chain amplification was performed using the Multiplex Fluorescent V2 kit (Biotechne, 323110) with the following conditions, with a 2 x 2 min wash between each step:

1. Amplifier 1, 30 min at 40°C
2. Amplifier 2, 15 min at 40°C
3. Amplifier 3, 30 min at 40°C
4. Channel 1 specific:

a. Amplifier 4, 15 min at 40°C
b. Custom Amplifier 5, 30 min at room temperature
c. Custom Amplifier 5, 15 min at room temperature
d. TSA Biotin (Akoya, NEL749A001KT) deposition, 15 min at room temperature
5. Hydrogen peroxidase block, 15 min at 40°C
6. Channel 2 specific:

a. Amplifier 4, 15 min at 40°C
b. Custom Amplifier 5, 30 min at room temperature
c. Custom Amplifier 5, 15 min at room temperature
d. TSA DIG (Akoya, NEL748001KT) deposition, 15 min at room temperature

The slides were washed for 2 x 5 min with MIBI Wash Buffer (1X TBS-T, 0.1% BSA), then subsequently blocked in Antibody Blocking Buffer (1X TBS-T, 2% Donkey Serum, 0.1% Triton X-100, 0.05% Sodium Azide) for 1 hour before the addition of the antibody cocktail (antibodies diluted in 1X TBS-T, 3% Donkey Serum, 0.05% Sodium Azide) overnight at 4°C. The following day, slides are washed for 3 x 10 min with MIBI Wash Buffer before a 15 min cross-linking with MIBI Crosslinking Buffer (1X PBS containing 4% PFA and 2% glutaraldehyde). Slides are then quenched briefly in 1X TBS-T, before being subjected to a series of washes and dehydration steps (3 x 100mM Tris pH 7.5, 3 x ddH2O, 1 x 70% EtOH, 1 x 80% EtOH, 2 × 95% EtOH, 3 x 100% EtOH).

For IF and MIBI cross validation PANINI experiments, sequential glass and gold slides containing both a 3D8 and 174xCEM pellet were processed exactly as described above, with the exception that the 2^*nd*^ hapten deposited was TSA PLUS Cy3 (Akoya, NEL744001KT). The glass slides also did not undergo a cross-linking step (which is a MIBI-specific processing step), but instead was subject to an anti-mouse secondary antibody 647 (Biolegend, Poly4053) for 1 hour before 3 x 10 min wash MIBI Wash Buffer, DAPI staining, cover-slipped and image processing on a Keyence BZ-X800 microscope with a Nikon CFI Plan Apo lambda 20x object (NA 0.75). In all PANINI experiments on gold slides containing SIV-positive and SIV-negative rhesus macaque lymph node tissue sections, glass slide controls containing sequential tissue sections and the 3D8/174xCEM cell pellets were ran in parallel.

### MIBI-TOF Data Acquisition and Processing

Mass images were acquired on a custom alpha-iteration MIBI-TOF mass spectrometer equipped with a duoplasmatron ion source (Ionpath) running research grade oxygen (Airgas, OX R80). All 196 multiplexed images in this study, an accumulation of 19404 individual channel TIFFs, were acquired using the following parameters:

Pixel dwell time: 12 ms

Image size: 400 um x 400 um at 512 x 512 pixels

Probe size: ~400 nm

Primary ion current: 3.5 nA as measured via a Faraday cup on the sample holder

Number of depths: 3

MIBI images were extracted and denoised using MIBIAnalysis tools (https://github.com/lkeren/MIBIAnalysis) as previously described (Keren et al., 2018). All three depths were aligned and summed for all downstream analysis. A detailed description of this algorithm can be found here (Baranski et al., 2021).

### Image Segmentation

Cell segmentation was performed using a local implementation of Mesmer, which utilizes the DeppCell library (deepcell-tf 0.6.0) as described (Greenwald et al., 2021; Valen et al., 2016). We adapted the included *multiplex_segmentation.py* python script from the deepcell-tf library and imported the neural network weights for prediction from https://deepcell-data.s3-us-west-1.amazonaws.com/model-weights/Multiplex_Segmentation_20200908_2_head.h5). The input for the segmentation were denoised MIBI images for dsDNA (for nuclear features) and CD45 (for membrane features). Signals from these images were capped at the 99.7th percentile. Utilization of model_mpp = 1.8 in the *multiplex_segmentation.py* script uniformly generated the most ideal segmentation results for all the FOVs in this study.

### MIBI Image Analysis and Cell Type Annotation

Features from single cells in segmented MIBI images were extracted based on the segmentation generated above and written out as FCS files. FCS fields are then uploaded onto CellEngine (Primity Bio) to visually assess data quality and concatenate FCS files that pass the visual check for the presence of dsDNA and Histone H3 nuclear markers. All subsequent analysis is done using R. While all samples were processed experimentally and computationally in parallel, we further ensured normalization of per FOV signal variation by normalization the markers for each cell on a per-FOV basis using the FOV-specific median Histone H3 levels. The data was then arcsinh transformed with a cofactor of 1, followed by a capping of the signal 99.9^*th*^ percentile. Finally, single-cell data was rescaled to a 0 - 1 range for each marker.

Unsupervised classification of cell types was then performed on this scaled data with FlowSOM (Gassen et al., 2015) and cell types were identified from each cluster with marker enrichment modeling (Diggins et al., 2017). The following markers were used for cell type identification: CD16, DC-SIGN, CD4, CD56, CD21, Pax-5, CD163, CD68, CD3, CD20, CD169, CD8a, CD11b, CD36, CD45, MPO, SMA, HLA-DR and CD138. Identified clusters were plotted in 2D space and carefully visually compared with the MIBI multiplexed images to confirm accuracy and specificity of the annotations. Clusters that did not meet the accuracy and specificity visual threshold were subject to further iterative clustering.

### Cellular Neighborhood Analysis

Cellular Neighborhoods were computationally defined using the 20 nearest neighbors (including self), followed by a k-means clustering of k = 11 as previously described (Schürch et al., 2020). The scripts for performing CN identification can be found at: https://github.com/nolanlab/NeighborhoodCoordination.

### Linear Discriminant Analysis

Each marker features, with the exception of vDNA and vRNA, were summed for the 20 nearest neighbors (including self) of each cell. These means of these summed marker features were calculated for each animal and CN within these animals. This resulted in 11 CNs from each of the 6 animals, for a total of 66 rows of data. This data was then subject to standardization to a mean of 0 and a variance of 1. Linear Discriminant Analysis was subsequently performed using the *Ida* function in the *MASS* R package, with the grouping set to the identifier of each individual animal.

### Marker Correlation Analysis

The Pearson correlations of marker expressions on cell types were calculated using the *rcorr* function of the Hmisc R package. The Euclidean distance between correlation coefficient values between markers were computed and hierarchically clustered using the *hclust* function of the stats R package.

### Cell Interaction Analysis

The Delaunay triangulation of cells were identified by their cartesian XY position within each field of view using default setting from the deldir R package. Interacting cells and their coordinates were extracted from the delsgs output of *deldir*, and the distances between cells joined together by the edge of a Delaunay triangle were calculated within the two-dimensional space according to the following formula:

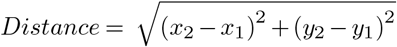

Cell to cell interactions within 100um from one another were identified, resulting in 1390517 interactions of the total 1392033 interactions observed between cells.

To establish a baseline distribution of distances, the same triangulation calculation was performed 1000 times, where for each iteration, the cell and neighborhood identified in each field of view were randomly assigned to existing XY positions. The average distance of a cell-cell interaction in each field of view for each permutation was calculated and this set of expected baseline distances was compared to the observed distances with a Wilcoxon Test.

The fold enrichment of distances between the observed data over the mean distances from the permutation test were calculated as follows:

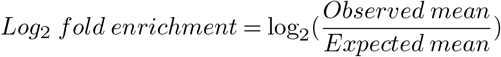

The log fold of the distances for each cell type and neighborhood interaction where p-values less than 0.05 were plotted for each group using ggplot2 in R.

### SIV-Infected Cells

All cells with a positive vDNA signal were marked as SIV-infected, before visually inspected to ascertain viral signal positivity and cell type annotation accuracy. Predominantly, SIV-infected cells were CD4 T cells, macrophages or Tregs. Rare cases of other cell types (such as B cells or endothelial cells) were deemed to be off-target effects and discarded from further investigation. These infected cells were then further divided into latent (vRNA = 0) or active (vRNA > 0) for the purpose of this study. Tregs were removed from all further analysis due to their small representation (n = 94) compared to the other 2 groups (CD4 T cells, n = 543; macrophages, n = 277).

### Random Forest Classification

A random forest classifier was used to examine if features of the tissue microenvironment could be used to identify latent and active SIV cells. Optimal parameters for the random forest model were identified using *trainControl* from the caret R package. Bootstrapping was implemented by randomly pulling 64% of the data as the training group and applying the classifier to the remaining 36% of the testing data to predict a cell’s reactivation status. The performance is reported as the median value from 100 repetitions and was evaluated by calculating the true positive rates, false positive rates, and the AUC of the resulting ROC as previously described (Robin et al., 2011). The predicted probabilities were then compared to the true reactivation status using a Wilcoxon test. Note that both vRNA and NFkB-p100 markers were removed from features used for the random forest classifier as they were molecular determinants of viral transcription.

### Binned Anchor Plot Analysis

A schematic of the anchor plot analysis is depicted in Figure 7C. All cells within a 100um range were extracted (1 pixel = 0.78125um), and the frequency of cell type and marker expression summed in 10um increments. These values were then divided by the number of cells, to normalize for differences in cell numbers in a radial spread from the center “anchor” cell. The 95% confidence interval for each binned value was then calculated and plotted along with the mean. Anchor plots were segregated by 1. Cell type (infected CD4 T cell or infected macrophage) and 2. Latent status (latent or active).

### Data Visualization

All pseudo-colored MIBI images were visualized using a Nolan lab specific instance of the MIBITracker (Ionpath). Figures 1A, 4A, 6E, 7C and 7G were generated in part using Biorender. All other plots in this manuscript were generated with the *ggplots2* R package (Wickham, 2016).

## Supplementary Figures

**Figure S1: Related to Figure 1.**
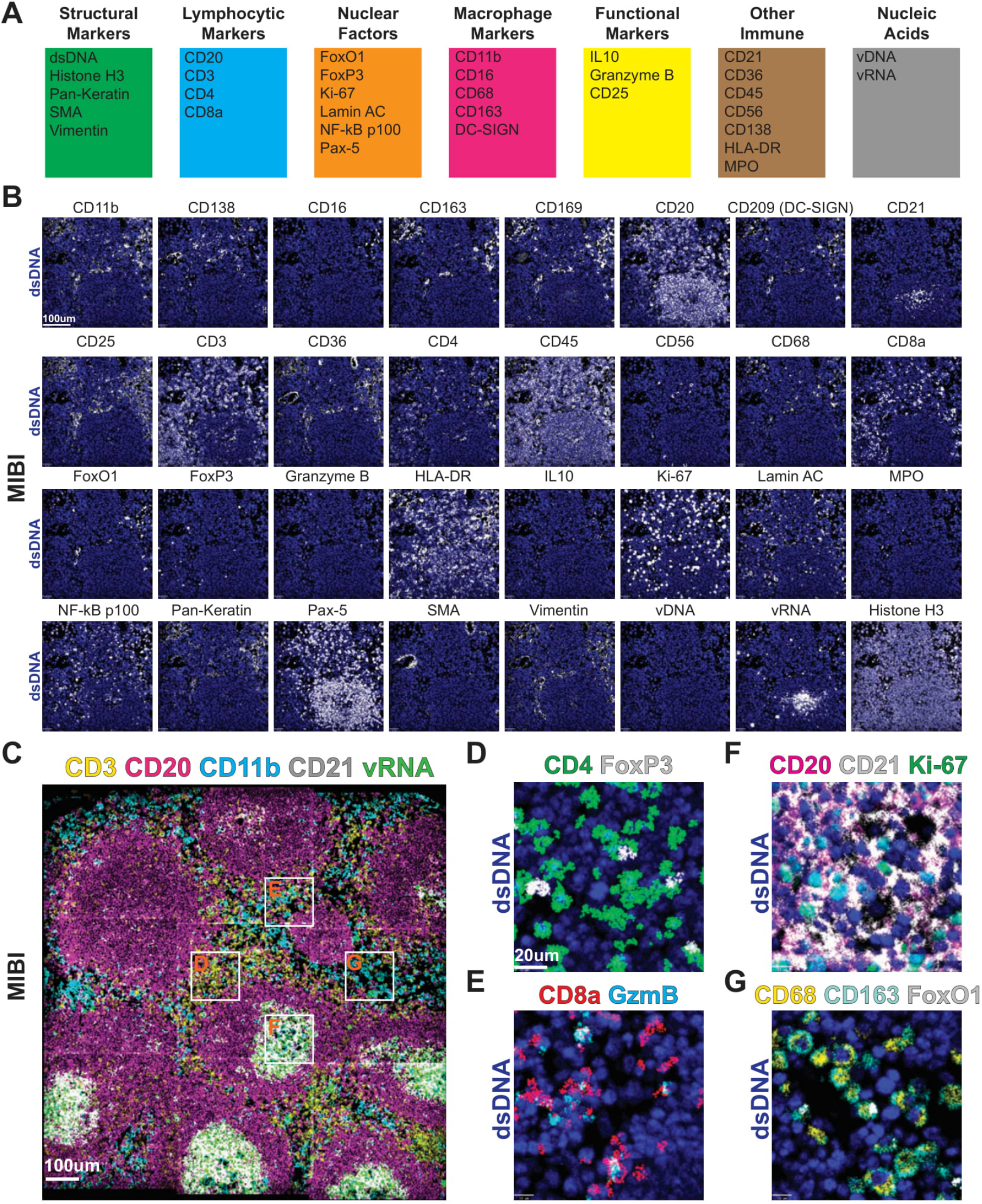
**(A)** The validated rhesus macaque compatible marker panel used in this study. **(B)** Images for each of the 33 markers depicted in pairwise fashion with dsDNA (blue). The FOV represented here is a germinal center within an SIV-positive lymph node. **(C)** A large FOV representing a 1.2 mm x 1.2 mm region of a SIV-positive lymph node with a number of lineage-specific markers. White boxes indicated regions magnified in the following panels. **(D)** A magnified region of panel C containing CD4- and FoxP3-positive T cells. **(E)** A magnified region of panel C containing CD8- and Granzyme B-positive T cells. **(F)** A magnified region of panel C containing CD20-, CD21- and Ki-67-positive B cells and FDCs. **(G)** A magnified region of panel C containing CD68-, CD163-, and FoxO1-positive macrophages.

**Figure S2: Related to Figure 2.**
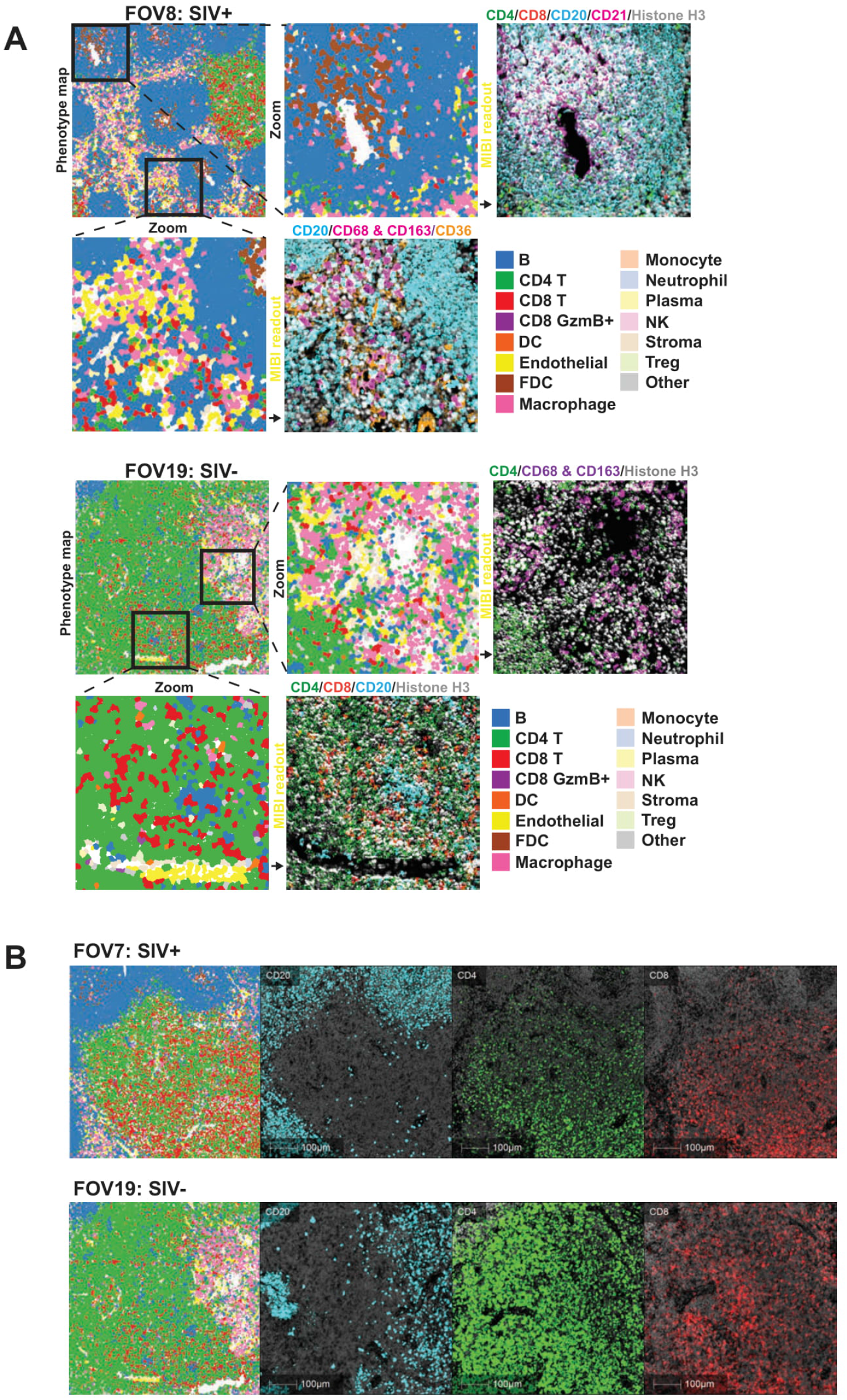
**(A)** Phenotype maps of two FOVs with two magnified regions of both the phenotype map and the paired pseudo-colored MIBI image with lineage-specific markers to validate the computationally determined cell phenotypes. **(B)** Representative FOVs from an SIV-positive and a control lymph node with three subjacent tissue sections that were stained for CD20 (blue), CD4 (green), and CD8 (red) and imaged using an IF microscope for orthogonal validation of PANINI-MIBI staining. The antibody clones and staining conditions used for the IF validation were identical to PANINI-MIBI.

**Figure S3: Related to Figure 2.**
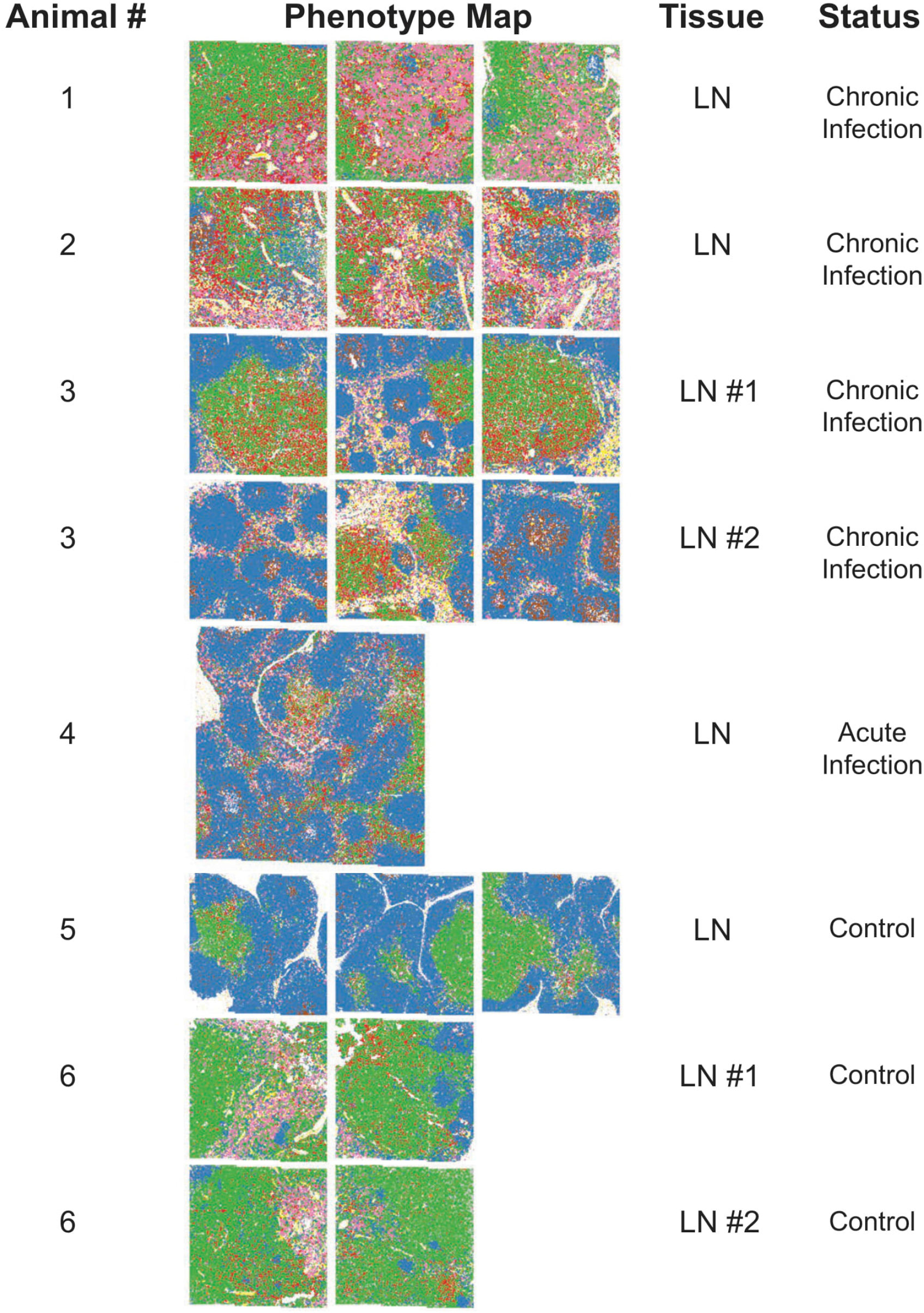
Phenotype maps of all 20 FOVs and their associated tissue sources. All FOVs are 1.2 mm x 1.2 mm with the exception of that from Animal 4 (2 mm x 2 mm).

**Figure S4: Related to Figure 3.**
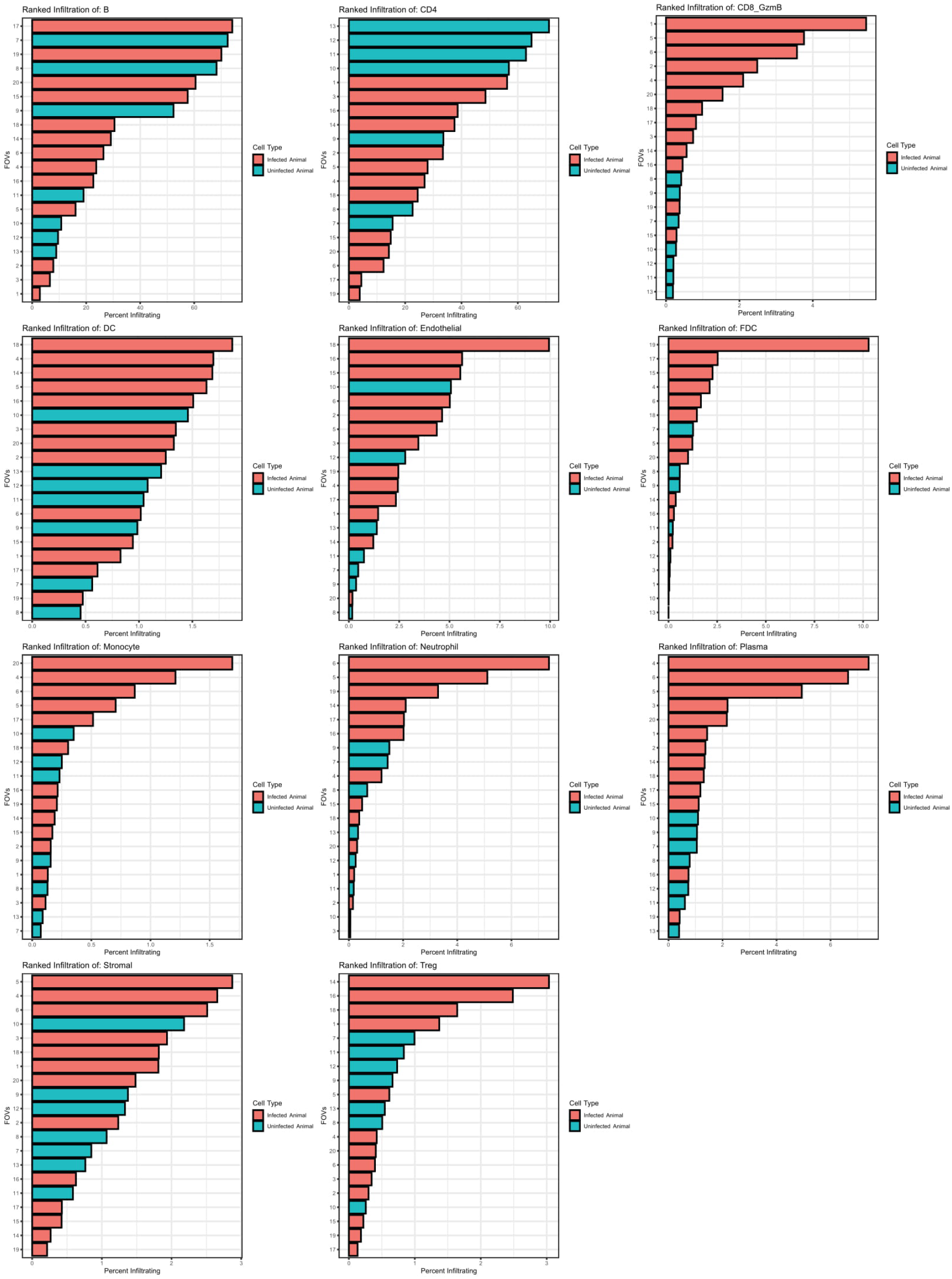
Ranked bar plots showing the percent infiltration of each cell type for the 20 FOVs with bars colored by infection status.

**Figure S5: Related to Figure 4.**
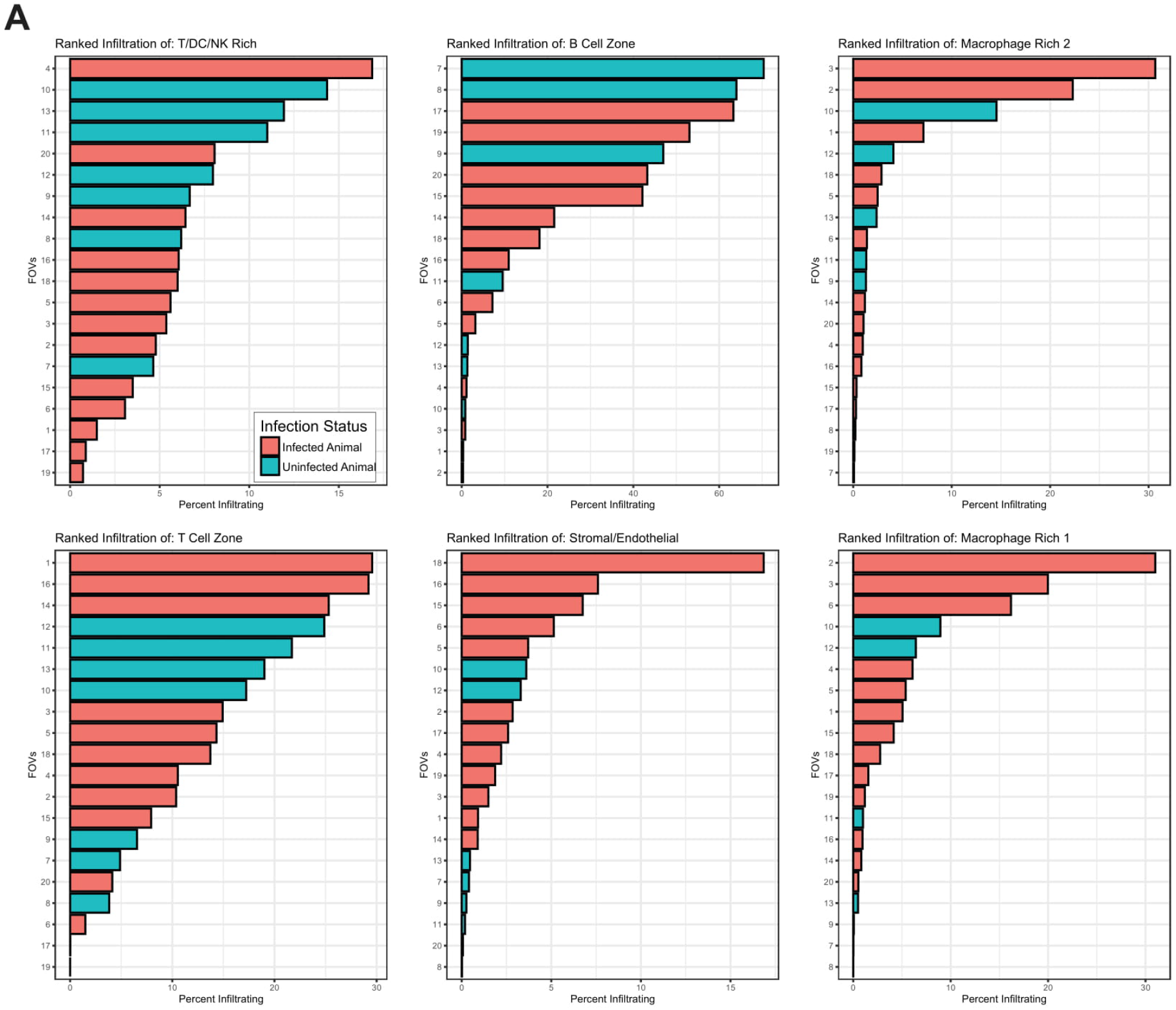
**(A)** Ranked bar plots showing the percent infiltration of each CN across the 20 FOVs with bars colored by infection status.

**Figure S5: Related to Figure 4.**
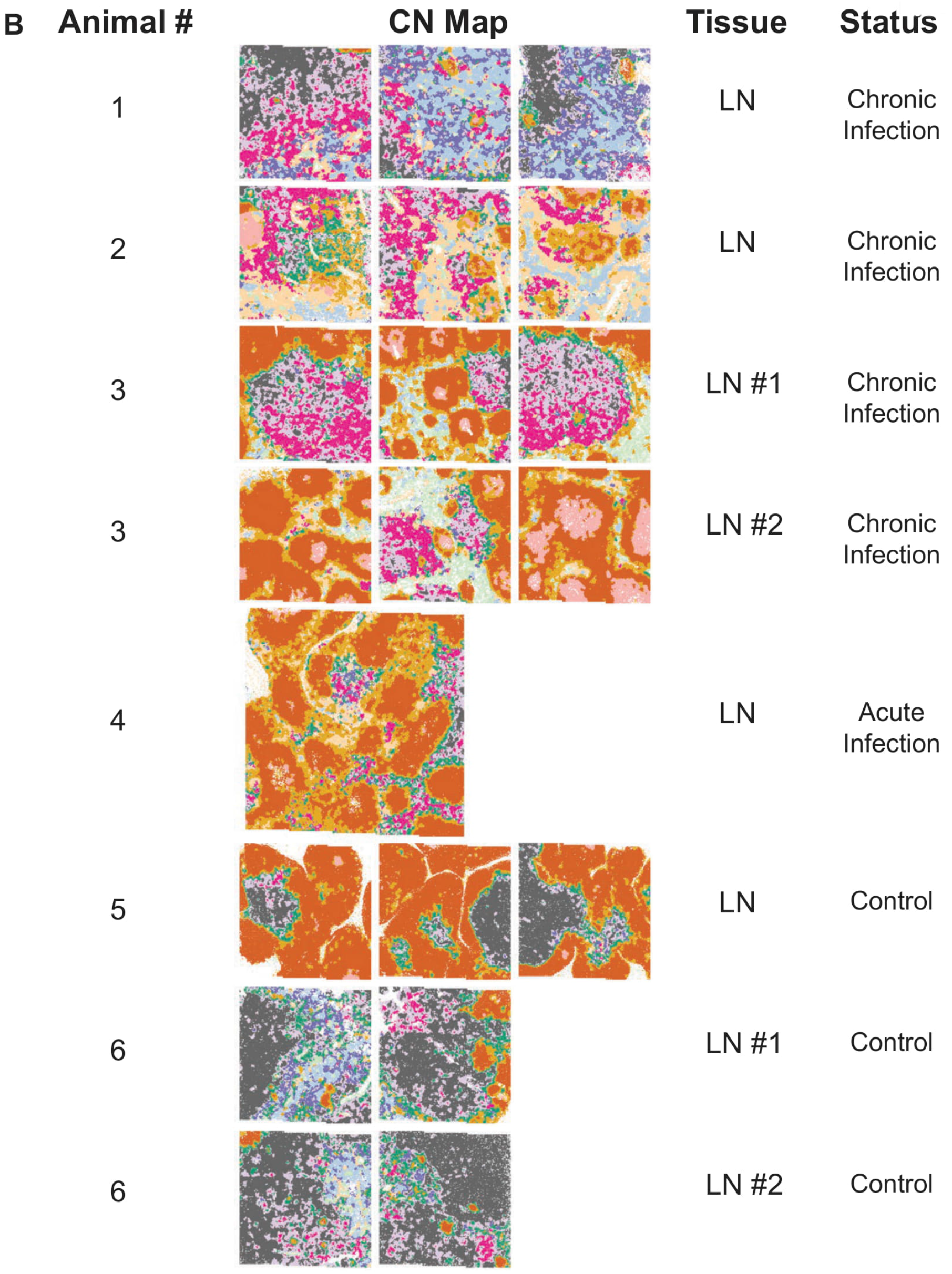
**(B)** CN maps of all 20 FOVs and their associated tissue sources. All FOVs are 1.2 mm x 1.2 mm with the exception of that from Animal 4 (2 mm x 2 mm).

**Figure S6: Related to Figure 4.**
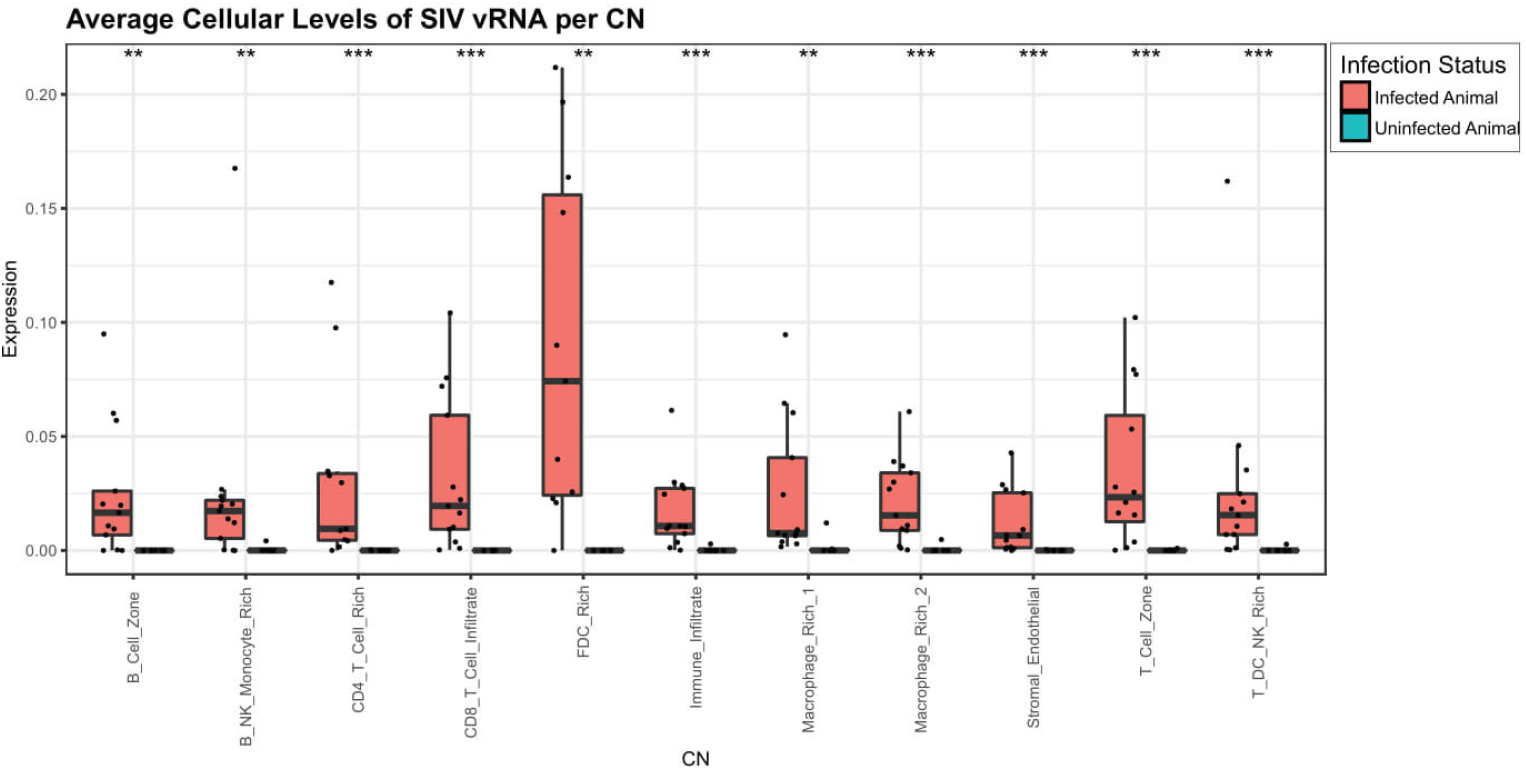
The mean SIV vRNA levels per CN. Each dot represents an individual FOV from an infected (orange) or uninfected (teal) animal.

**Figure S7: Related to Figure 5.**
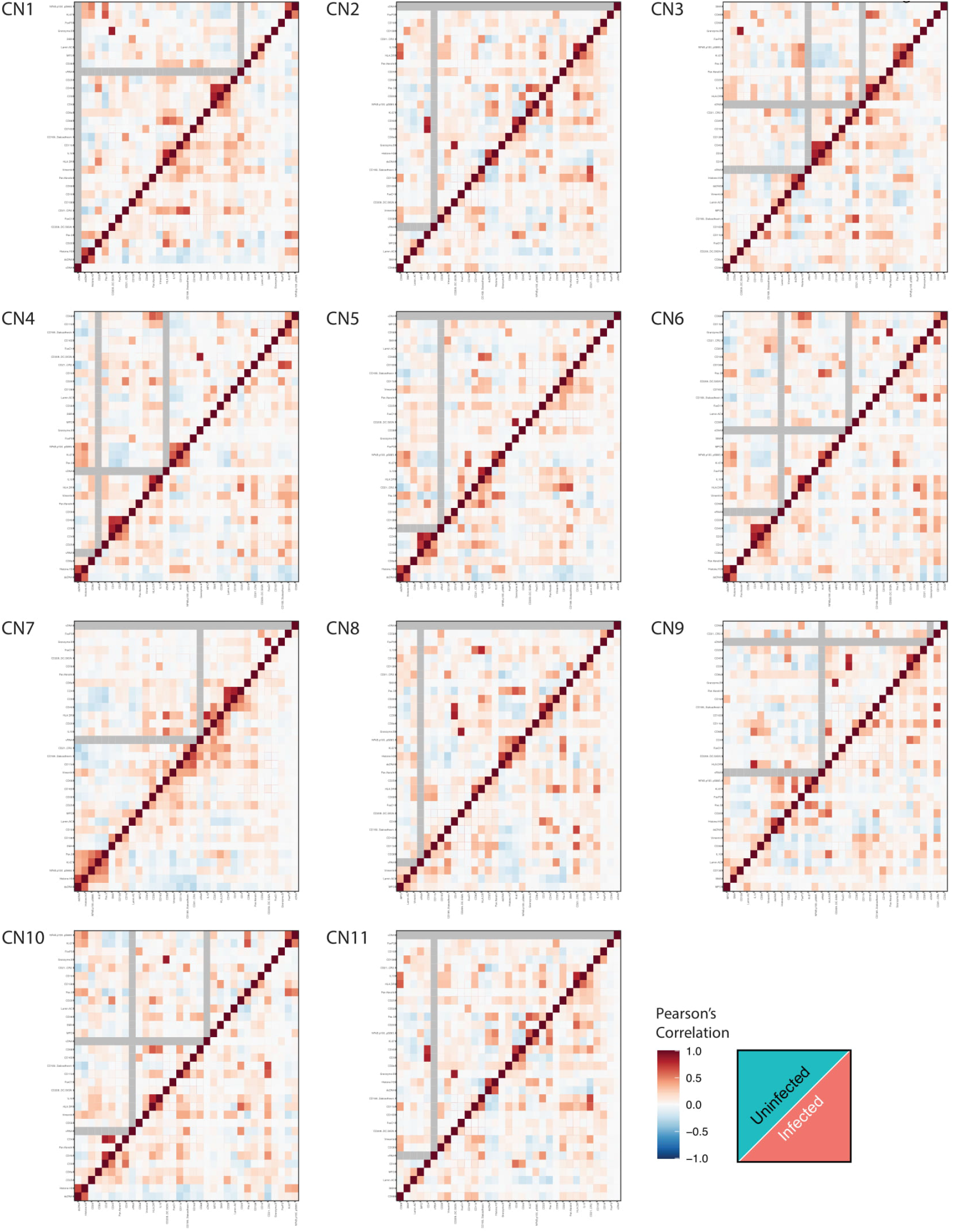
Heatmaps of pairwise Pearson’s correlations of markers across each individual cell within each CN for infected (top left) and healthy (bottom right) animals.

**Figure S8: Related to Figure 7.**
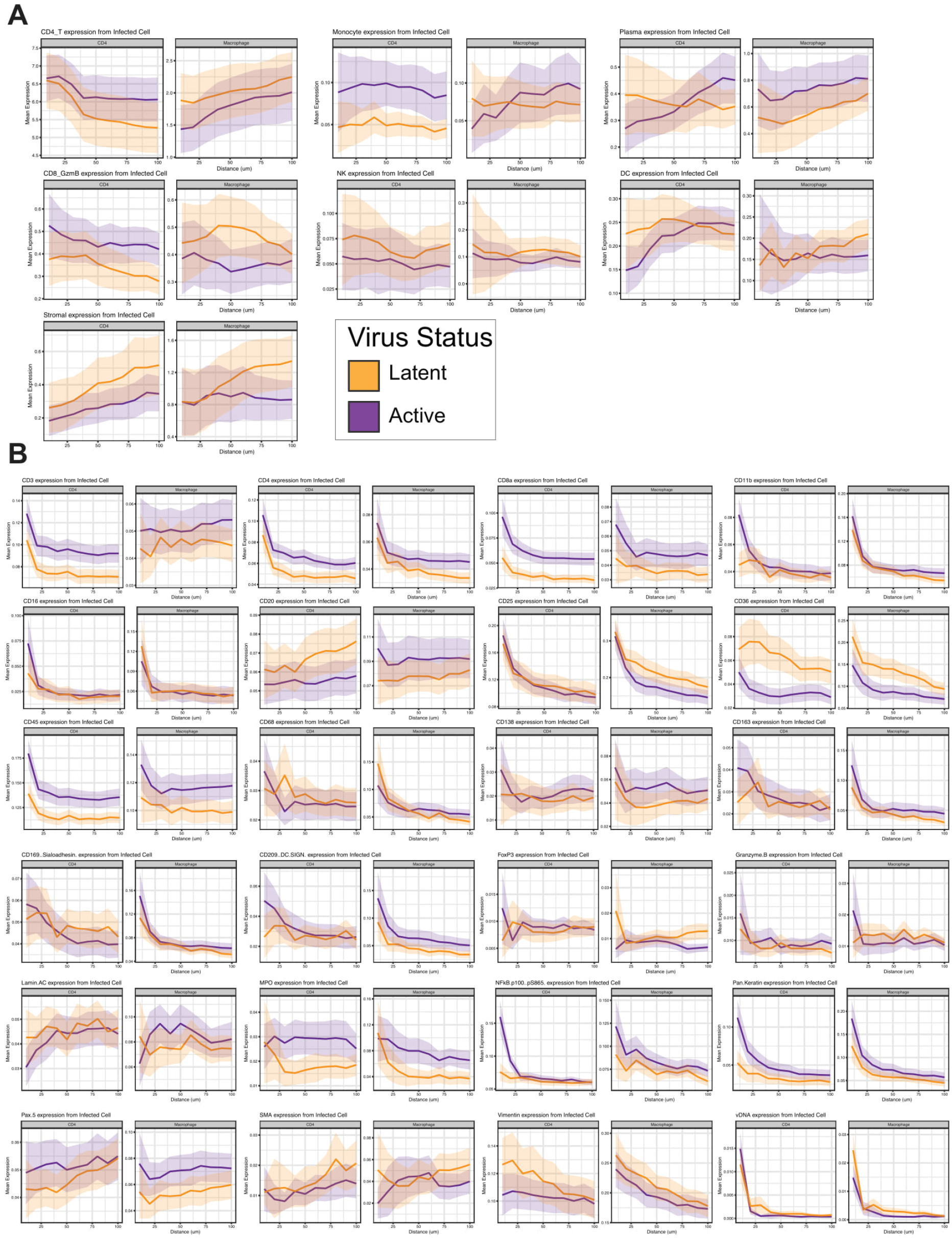
**(A and B)** Anchor plots of A) mean cell type quantifications and B) mean marker expression around infected CD4^+^ T cells (top) or macrophages (bottom). Orange indicates latent cells, and purple indicates transcriptionally active cells. The thick colored lines represent the mean values, and the light regions around these lines depict the 95% confidence intervals. The infected cells were anchored at 0 μm, and the plot ends at 100 μm.

