## Supplementary material for "Virus-Dependent Immune Conditioning of Tissue Microenvironments": Key Resource Table

### KEY RESOURCES TABLE

| REAGENT or RESOURCE | SOURCE | IDENTIFIER |
|---|---|---|
| **Antibodies** | | |
| dsDNA (Clone 35I9 DNA) | Abcam | ab27156 |
| Vimentin (Clone D21H3) | Cell Signaling Technology | Custom Order for Carrier Free |
| Histone H3 (Clone D1H2) | Cell Signaling Technology | Custom Order for Carrier Free |
| CD16 (Clone D1N9L) | Cell Signaling Technology | Custom Order for Carrier Free |
| SMA (Clone D4K9N) | Cell Signaling Technology | Custom Order for Carrier Free |
| CD209 (DC-SIGN) (Clone DCN46) | Biolegend | 551186 |
| NFkB-p100 (pS865) (Polyclonal) | Abcam | ab31474 |
| CD4 (Clone EPR6855) | Abcam | ab181724 |
| CD56 (Clone MRQ-42) | Cell Marque | Custom Order for Carrier Free |
| FoxP3 (Clone 236A/E7) | Thermo Fischer Scientific | 14-4777-82 |
| Granzyme B (Clone EPR20129-217) | Abcam | ab219803 |
| CD21 (CR2) (Clone SP186) | Abcam | ab240987 |
| Ki-67 (Clone 8D5) | Cell Signaling Technology | Custom Order for Carrier Free |
| Pax-5 (Clone D7H5X) | Cell Signaling Technology | Custom Order for Carrier Free |
| CD138 (Clone EPR6454) | Abcam | ab226108 |
| CD163 (Clone EDHu-1) | Novus | NB110-40686 |
| CD68 (Clone D4B9C) | Cell Signaling Technology | Custom Order for Carrier Free |
| FoxO1 (Clone C29H4) | Cell Signaling Technology | Custom Order for Carrier Free |
| CD3 (Clone MRQ-39) | Cell Marque | Custom Order for Carrier Free |
| CD20 (Clone SP32) | Abcam | ab64088 |
| Lamin A+C (Clone EPR4100) | Abcam | ab216074 |
| MPO (Polyclonal) | R&D Systems | AF3667 |
| HLA-DR (Clone EPR3692) | Abcam | ab215985 |
| IL10 (Clone 4A7-25-17) | Abcam | ab134742 |
| CD8a (Clone D8A8Y) | Cell Signaling Technology | Custom Order for Carrier Free |

| | | |
|---|---|---|
| Pan-Keratin (Clone AE1/AE3) | Biolegend | 914204 |
| CD11b (Clone EPR1344) | Abcam | ab209970 |
| CD36 (Clone D8L9T) | Cell Signaling Technology | Custom Order for Carrier Free |
| CD25 (Clone 4C9) | Cell Marque | 125M-16 |
| CD45 (Clone D9M8I) | Cell Signaling Technology | Custom Order for Carrier Free |
| Anti-Biotin (Clone 1D4-C5) | Biolegend | 409002 |
| Anti-Digoxigenin (Clone 21H8) | Abcam | ab420 |
| **Bacterial and Virus Strains** | | |
| SIVmac251 | AIDS reagent resource | 253 |
| SIVmac239 | Mudd et al 2018. PMID: 30262807 | N/A |
| **Biological Samples** | | |
| FFPE Inguinal LN from acute SIVmac239 infected rhesus macaque (Day 13 p.i.) | Mudd et al 2018. PMID: 30262807 | RHCF4T |
| FFPE Mesenteric LN from chronic SIVmac239X infected rhesus macaque (week 16 p.i.) | Oregon National Primate Research Laboratory | 34675 |
| FFPE Mesenteric LN from chronic SIVmac239X infected rhesus macaque (week 19 p.i.) | Oregon National Primate Research Laboratory | 34622 |
| FFPE Mesenteric LN from chronic SIVmac251 infected rhesus macaque (Day 227 p.i.) | Oregon National Primate Research Laboratory | 33098 |
| FFPE Mesenteric LN from SIV neg rhesus macaque | Oregon National Primate Research Laboratory | 32518 |
| FFPE Inguinal LN from SIV neg rhesus macaque | NCI/ACVP | A7E033A |
| **Chemicals, Peptides, and Recombinant Proteins** | | |
| TBS IHC Wash Buffer plus Tween 20 | Sigma | 935B-09 |
| Dako Target Retrieval Solution, pH 9 | Agilent | S236784-2 |
| Dako Target Retrieval Solution, pH 9 | Agilent | S236784-2 |
| Avidin/Biotin Blocking Kit | Biolegend | 927301 |
| Glutaraldehyde 10% Aqueous Solution EM Grade | EMS | 16120 |
| Donkey Serum | Sigma | D9663-10ML |
| 16% Paraformaldehyde (formaldehyde) aqueous solution | EMS | 15711 |
| VECTABOND Reagent for Tissue Section Adhesion | Vector Labs | SP-1800 |
| TCEP | Sigma | C4706-10G |
| Candor PBS antibody stabilizer | Fisher Scientific | NC0436689 |
| **Critical Commercial Assays** | | |
| RNAscope Multiplex Fluorescent Detection Kit V2 | Biotechne | 323110 |
| RNAscope v2.5 HD Detection Brown | Biotechne | 322310 |
| SIVmac239-gag-pol sense (vDNA) | Biotechne | 416141 |

| | | |
|---|---|---|
| SIVmac239-vif-env-nef-tar (vRNA) | Biotechne | 416131-C2 |
| Maxpar X8 Multimetal Labeling Kit | Fluidigm | 201300 |
| Ionpath Conjugation Kits | Ionpath | 600XXX |
| TSA Plus Biotin 50-150 slides | Akoya | NEL749A001KT |
| TSA Plus DIG, 50-150 Slides | Akoya | NEL748001KT |
| **Deposited Data** | | |
| Multiplexed Images | This Study | Data available at: www.mibi-share.ionpath.com |
| **Experimental Models: Cell Lines** | | |
| 3D8 | AIDS reagent resource | 13239 |
| 174XCEM | AIDS reagent resource | 272 |
| **Experimental Models: Organisms/Strains** | | |
| Rhesus macaques of Indian origin | ONPRC/NCI | N/A |
| **Oligonucleotides** | | |
| N/A | | |
| **Recombinant DNA** | | |
| N/A | | |
| **Software and Algorithms** | | |
| Matlab 2019b | Mathworks | |
| MIBIAnalysis | Keren et al 2018 | https://github.com/lkeren/MIBIAnalysis |
| R 3.6.3 | | https://www.r-project.org/ |
| Cellular Neighborhoods | Schürch et al 2020 | https://github.com/nolanlab/NeighborhoodCoordination |
| DeepCell 0.6.0 | Greenwald et al 2021 | https://github.com/vanvalenlab/deepcell-tf |
| Mesmer | Greenwald et al 2021 | https://github.com/vanvalenlab/deepcell-tf |
| CellEngine | Primity Bio | https://cellengine.com/ |
| FlowSOM | Van Gassen et al 2015 | https://bioconductor.org/packages/release/bioc/html/FlowSOM.html |
| MEM | Diggins et al 2017 | https://github.com/cytolab/mem |
| Hmisc R package | N/A | https://cran.r-project.org/web/packages/Hmisc/index.html |
| MASS R package | Venables WN, Ripley BD (2002) | https://cran.r-project.org/web/packages/MASS/index.html |

| | | |
|---|---|---|
| deldir R package | N/A | https://cran.r-project.org/web/packages/deldir/index.html |
| caret R package | N/A | https://cran.r-project.org/web/packages/caret/index.html |
| ggplots2 R package | N/A | https://cran.r-project.org/web/packages/ggplot2/index.html |
| MIBITracker | N/A | https://mibi-share.ionpath.com/ |
| **Other** | | |
| MIBIscope (Alpha Iteration) | Ionpath | N/A |
| ACD HybEZ Hybridization System | Biotechne | 310013 |
| Lab Vision PT Module | Fisher Scientific | A80400012 |
